# Cooperative reading of DECA-CCAAT composite element by the TALE/NF-Y/Sp2 transcription factors

**DOI:** 10.64898/2026.02.16.706105

**Authors:** Andrea Bernardini, Nerina Gnesutta, Roberto Mantovani

## Abstract

To understand how genes are regulated it is important to assess the molecular basis of TFs cooperation in promoter and enhancer regulatory elements. Compelling genetic, genomic and biochemical data in mammals and zebrafish point at DECA-CCAAT composite elements as important for regulation of gene expression in development. DECA is recognized by the homeodomain (HD) heterodimeric TALE TFs (PBX/PREP), CCAAT by the NF-Y trimer. Both are evolutionarily conserved in eukaryotes, including plants. Sp2, a member of the Sp/KLF family, potentiates association of TALE and NF-Y on DNA. We applied AlphaFold to infer the structural basis of the hexameric complex with DNA. The resulting models position TALE and NF-Y on the respective sites, predicting the complex arrangement of the MEINOX/PBC heterodimerization module of the TALE TFs and the sequence-specific DNA contacts of the HDs to DECA. The complex is tied by the Sp2 N-terminus, which contacts the TALE dimer through the α-helical SP-box and the NF-Y trimer through a novel <u>s</u>hort <u>li</u>near <u>m</u>otif (YA-SLiM). A flexible linker separates the anchor points, consistent with the constrained stereo-alignment of the DECA-CCAAT motif. The models are confirmed by assembly of the recombinant NF-Y/TALE/Sp2 in complex with DNA *in vitro.* Mutagenesis confirms the importance of the SP-box and YA-SLiM for cooperativity. Rationalising available genomic and biochemical data, the structural models depict a novel mode of cooperative binding on DNA, providing important clues as to how TFs potentially function beyond DECA-CCAAT in different eukaryotic contexts.

## INTRODUCTION

Transcriptional regulation is key for development, differentiation and response to physiological and pathological stimuli of eukaryotes. A fundamental aspect is initiation of transcription, which depends on binding of Transcription Factors-TFs-to discrete DNA sequences in promoters and enhancers. TFs are arranged in a precise and coherent way within regulatory regions (1). The human genome has some 1600 genes coding for TFs, typically classified according to the DNA-binding domains-DBD-used to select the DNA elements (2). A large family is represented by homeodomains-HD-in turn classified in several sub-families (3). TF binding sites are aligned in promoters with a precise, evolutionarily conserved logic and a key issue is understanding their functional synergism. Multiple mechanisms were shown, including cooperative DNA-binding: the affinity of one TF for its site might be suboptimal, or indeed weak, but in the presence of neighbouring TFs, stable complexes with DNA are formed. Structural studies on the β-Interferon promoter provided a paradigm as to the precise positioning of multiple TFs required for full promoter activity (4).

Together with the related MEIS1-3 (Myeloid Ecotropic viral Insertion Site) (5), PREP1/2 (PBX-regulating protein), also known as PKNOX1/2 (PBX/Knotted homeobox) according to the plant nomenclature, belong to the TALE (Three Amino acids Loop Extension) sub-family of homeodomain (HD) TFs (6). They are named after an extension of three amino acids between helix 1 and 2 of the three α-helices of the HD that mediate DNA-binding. They heterodimerize with PBX1-4 (Pre-B-cell leukaemia homeobox) partners, also harbouring a HD (7): formation of the heterodimer allows nuclear localization (8), efficient DNA-binding through the respective HDs to an asymmetric decameric motif (DECA) and transcriptional activation (9, 10).

The N-terminal half of PREP contains the ancestral MEINOX domain shared by mammals and plants (5, 11), which mediates heterodimerization, as assessed by deletion or site directed mutagenesis, which further pinpointed two subdomains, termed HR1/2, necessary for heterodimerization and transfer of the complex to the nucleus (7, 8, 12, 13). HR1/2 are predicted to be α-helical, each with two heptad leucine/isoleucine-rich motifs, remindful of leucine zippers (14). PBX association with the TALE partners is mediated by the PBC domain (divided into PBC-A and PBC-B) also located at the N-terminal half of the protein and predicted to form α-helices (15). The affinities for the partners are apparently different, with PREP>MEIS; the two TFs compete for PBX binding when expressed in the same cell line, often with opposing outcomes: PREP has been generally associated to a tumour-suppressor role, MEIS to an oncogenic one (16, 17).

Genomic analyses found that PREP and MEIS share some locations, but mostly have independent sites, cobound by PBX1 (18, 19): in particular, PREP is mainly associated to promoters, MEIS to enhancers. In these studies, canonical TALE locations are often juxtaposed to CCAAT boxes, typical of eukaryotic promoters. PBX1/2 locations in mouse epiblast stem cells were also found enriched in CCAAT (20). The CCAAT element is bound by the NF-Y (CBF) TF, crucial in maintaining the upstream border of the core promoter free of nucleosomes (21) and whose removal causes relocation of transcription start site (TSS) and generation of aberrant transcripts. ChIP-seq and the use of Dominant Negative NF-YA mutant in zebrafish returned genes coregulated by NF-Y and PBX4 at Zygotic Genome Activation (3.5 hpf) (22, 23). Indeed, our latest analysis of ENCODE ChIP-seq data found PREP, MEIS2 and PBX2/3 among the TFs binding next to NF-Y sites with high frequency (5-20% of sites) in different cell lines (24). Interactions between NF-YA/NF-YB and PREP were reported with co-IP assays in zebrafish (22).

A third TF considered here is Sp2, a member of the large Sp/KLF family of TFs, which harbour a C2H2-Zinc Finger (ZF) DBD and a QVIT-rich Trans-Activation Domain (TAD) (25). Sp2 location analysis in mouse *wt* and *Sp2*^-/-^ embryonic fibroblasts found sites with multiple CCAAT boxes, rather than the predicted GC-rich matrix of Sp/KLFs (26, 27). Subsequently, these Authors reported that Sp2 associates to chromatin and functions through a DECA-CCAAT geometry: GAnnGAC-11bp-CCAAT. Sp2 appears not to bind DNA directly, since the ZF DBD was dispensable for chromatin association, but Sp2 facilitates DNA binding of the TALE/NF-Y complex by protein-protein interactions through its most N-terminal region (aa 1-94) (28). Precisely the same arrangement of sites was found in the zebrafish NF-Y/PREP/PBX4 coregulated units (22, 23). Thus, the TALE/NF-Y/Sp2 connection is evolutionarily conserved.

The experimental 3D structures of the yeast, mammals and plant NF-Y in complex with CCAAT are available (29–31). NF-YB/NF-YC Histone Fold Domains-HFD-contact the phosphate backbone of the DNA through basic residues of the L1/IZ1/L2 structures, similarly to the H2A/H2B heterodimer in nucleosomes (32). Sequence-specificity is provided by the NF-YA A2 helix and GXGGRF motif, contacting CCAAT nucleobases in the minor groove; in turn, this drives severe bending of the DNA-some 80°-which is stabilized by the flanking HFD L1/L2 pairs. Overall, there are >35 DNA contacts covering some 30 bp, an unusually long stretch for a sequence specific TF. Regarding TALE TFs, an experimental structure for the PREP1/PBX1 heterodimer is currently not available. There is 3D knowledge of the homeodomains (HDs), either in isolation or in complex with DNA: PBX1 in complex with HOXA9 or HOXB1, MEIS1 with HOXB13 or DLX3 (33–36). As for Sp2, no experimental structures are currently available.

AlphaFold 3 (AF3) is a powerful tool to predict macromolecular structures, being increasingly used to reconstruct binding of TFs to DNA (1, 37, 38). We decided to employ it to infer notions on DNA-bound PBX1/PREP1/NF-Y/Sp2/DNA complex. The exercise results in 3D models that are consistent with the available biochemical, structural -and genomic-data; they confirm and extend our knowledge on NF-Y, returning relevant information on the novel and complex fold of MEINOX/PBC heterodimerization module and on the DNA-binding mode of the TALE TFs. Interestingly, the model predicts the molecular basis of Sp2 tethering on the DNA-bound TALE/NF-Y platform through two short motifs, including the elusive SP-box and a novel conserved motif. The models explain the rules of distance between the sites, providing a novel paradigm about TF-TF cooperativity on DNA. Finally, we validate the models by employing recombinant proteins in DNA-binding assays.

## MATERIALS AND METHODS

### Remapping of available ChIP-seq data

To assess chromatin binding of Sp2, NF-Y subunits and PBX1/PREP1 and generate **Fig. 1B**, we remapped available ChIP-seq datasets from (27, 28). Raw fastq run files were downloaded from ENA database (see Data Availability section for accession numbers). Reads were aligned to the mouse genome (mm10) using bowtie2 (39) using the following options:--no-mixed--no-discordant-k 1. SAM files were converted to BAM files, sorted and indexed using SAMtools (40). BigWig files were generated using deepTools (41) using the command bamCoverage with the options--normalizeUsing RPKM--extendReads 200. ChIP-seq coverage tracks were inspected and visualised in IGV (42).

**Figure 1.**
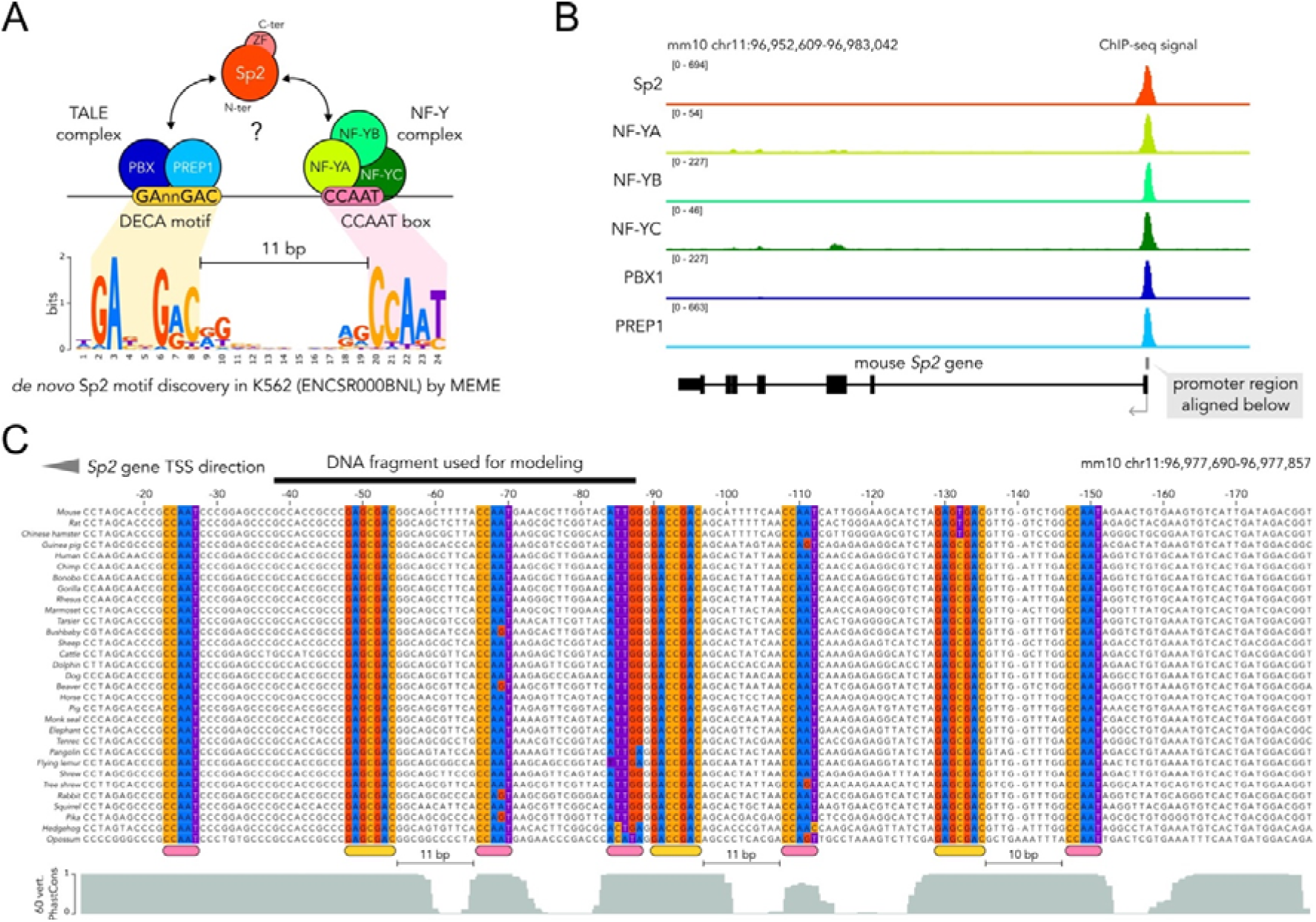
Overview of Sp2 genomic binding mode. **A**. Scheme depicting Sp2 genomic sites constituted by DECA-CCAAT sites bound by TALE (PBX/PREP) and NF-Y TFs. Sp2 binds these regions using its N-terminal domain (aa 1-94), while the ZF DBD is dispensable. The typical 11 bp spacing between DECA and CCAAT motifs bound by Sp2 is indicated above the sequence logo, obtained from *de novo* motif discovery of Sp2-bound regions in K562 cells by MEME. **B**. ChIP-seq signals showing binding of Sp2, NF-Y subunits, PBX1 and PREP1 on a representative Sp2 target (mouse Sp2 promoter). ChIP-seq experiments were performed in mouse embryonic fibroblasts (27, 28). **C**. Multi-species sequence alignment of the Sp2 promoter region shown in B (Multiz35Align). Positions with CCAAT and DECA motifs (pink and orange boxes below the alignment) are highlighted with the same nucleotide colour code used in A. The spacing between DECA and CCAAT motifs and the DNA region selected for structural modelling are indicated. The 60 vertebrate PhastCons conservation scores from UCSC genome browser are shown below the alignment.

### Structural modelling and visualisation

#### Structure prediction with AlphaFold

AlphaFold 3 (43) structural models were generated using the web server application (https://alphafoldserver.com/). No seed option was used. The AlphaFold2-multimer structural models were generated using ColabFold v1.5.5 Google Colaboratory (44) with default parameters. Intermolecular interface mapping plot shown in **Fig. 2B** was generated with AlphaBridge (45) using default confidence level for interface detection. PAE heatmaps were generated by uploading JSON output files in PAE Figure Editor (https://thecodingbiologist.com/tools/pae.html). Protein and DNA sequences used as input for AF modelling are listed in **Supplementary Table 1**. A list of the models generated and their description is documented in **Supplementary Table 2**.

**Figure 2.**
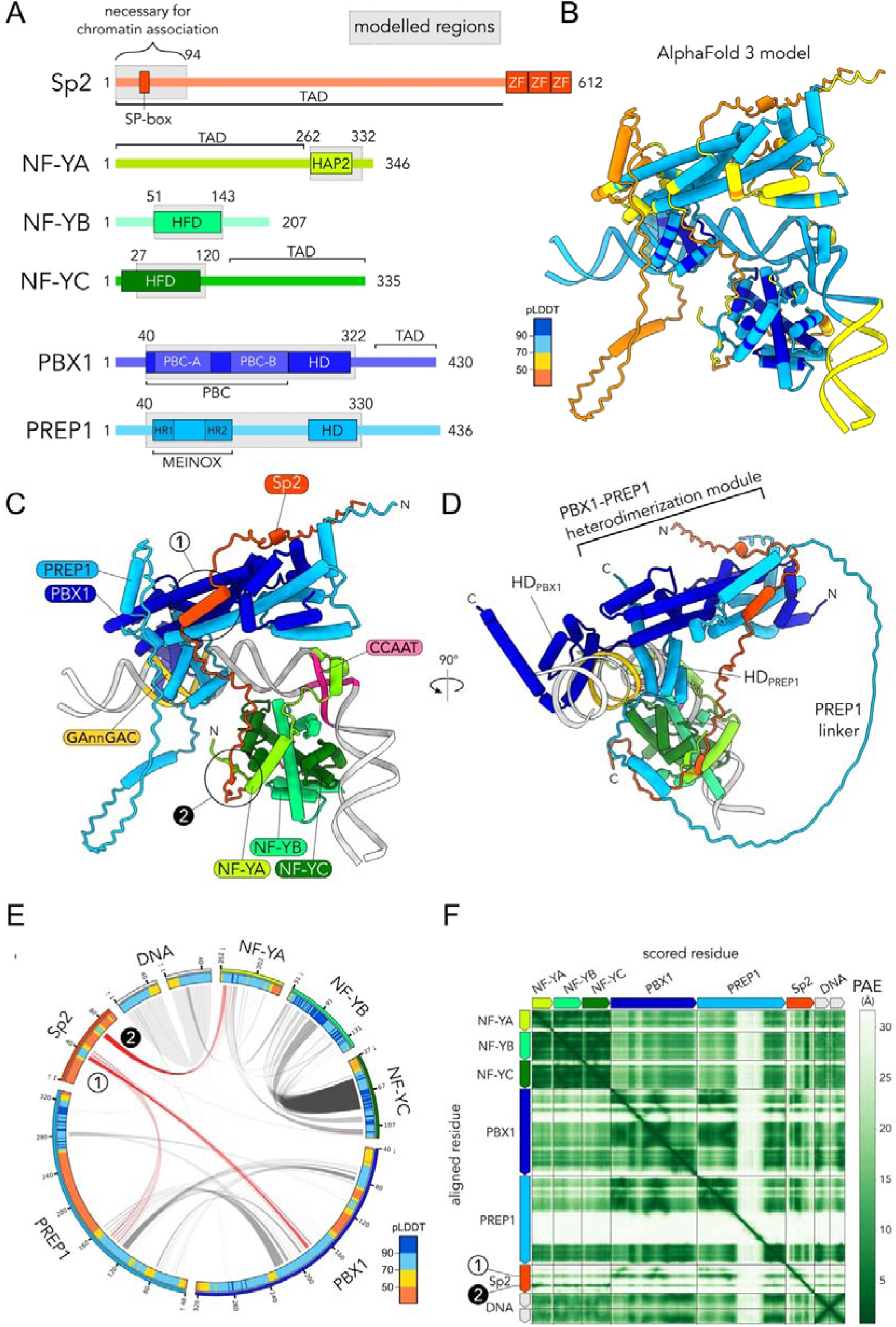
Overview of the minimal TALE/NF-Y/Sp2 complex on DECA-CCAAT DNA modelled by AlphaFold 3. **A**. Scheme of the mouse full length proteins considered here. The main structured domains of each protein are shown in coloured boxes, while characterized activation domains (AD) are indicated with squared parenthesis. The regions used to build models of the minimal-domain complex with AF3 are indicated with grey boxes. **B**. TALE/NF-Y/Sp2/DNA AF3 model coloured according to pLDDT confidence score colour scheme. **C**. AF3 model shown in B, coloured to distinguish each protein chain. DNA segments corresponding to the DECA and CCAAT motifs are coloured in yellow and pink, respectively. The two Sp2 anchor points contacting PBX1/PREP1 (#1) and NF-Y (#2) are circled. **D**. Rotated view of the model shown in C. **E**. Inter-molecular interface network of a representative TALE/NF-Y/Sp2/DNA AF3 model computed with AlphaBridge. Each molecular chain in the complex is shown along the circumference and coloured according to AF pLDDT confidence score. Arcs connecting different subunits represent three-dimensional binary interfaces in the predicted complex. Interactions of the two anchor points that connect Sp2 to PBX1/PREP1 (#1) and NF-Y (#2) are highlighted in red. **F**. Predicted aligned error (PAE) heatmap for the AF model shown in B. Darker green shades represent residue pairs spatially positioned with higher confidence within the complex.

#### Structural visualization and analyses

All molecular models were inspected and analysed using UCSF ChimeraX (v1.9) (46). H-bonds and contacts mapping in the structures were predicted and visualized in ChimeraX using the hbonds and contacts commands with default parameters. Protein contacts with DNA nucleobases were first mapped by uploading AlphaFold 3 or experimental structures on DNAproDB (47) and selecting Maj. Groove Min. Groove and Base as DNA moieties to be considered. Each mapped contact was then manually inspected in ChimeraX using the hbonds and contacts commands as above, noting protein side-chains involved in base recognition but modelled with low confidence (pLDDT<70). Structural model superimposition was performed using the matchmaker command with default settings in ChimeraX. To reproduce AlphaFold pLDDT color scheme, we used the following command in ChimeraX: color bfactor palette 0, darkorange:49.99, darkorange:50, yellow:69.99, yellow:70, deepskyblue:89.99, deepskyblue: 90, blue:100, blue.

#### Mapping cross-linking mass-spectrometry data on predicted structural models

We mapped cross-linking mass-spectrometry data from (13) on AlphaFold generated models. First, 10 models of the human PBX1/PREP1 heterodimer were generated using AlphaFold 3 with the protein sequences matching the experimental constructs (i.e. PBX1 aa 1-308, PREP1 aa 1-344). The resulting .cif files were converted to .pdb format using GEMMI tools (https://project-gemmi.github.io/wasm/) (48). The experimental BS3 cross-links were retrieved from the two experiments as reported in the supplementary material from (13) and are listed in **Supplementary Table 3**. The cross-links were mapped onto AF models using xlms-tools (49). The cross-linked positions were formatted in a text file compatible with xlms-tools and the tool was run on all 10 models with default settings, where Ca-Ca distance violation threshold is set >3.3 nm. The output was visualized in ChimeraX. All models resulted in the same cross-links fitting within the PBC/MEINOX heterodimerization module.

#### Extraction of per-residue average pLDDT confidence scores

AlphaFold 3 output contains per-atom pLDDT confidence scores. We computed an average per-residue pLDDT score to generate plots in **Fig. 6**. Each AlphaFold model was opened in ChimeraX and the pLDDT scores for the protein chain of interest were extracted using the command ‘info atoms / <CHAIN> attribute bfactor saveFile browse’. The resulting text files were processed with a custom R script using the tidyverse package (v2.0.0) (50). In brief, the scores in each model were grouped by residue and the mean per-residue pLDDT was calculated. The mean per-residue pLDDT score across models was also computed and plotted using ggplot2.

### Alignments and sequence analyses

Multiple sequence alignments were generated and visualized using Jalview (51). Evolutionary constrained regions analysis for Sp2 were derived from AMINODE (52). Structural disorder prediction of Sp2 was performed using Metapredict (53). AlphaMissense substitution pathogenicity scores were obtained from the AlphaFold Protein Structure Database (https://alphafold.ebi.ac.uk/).

### Expression constructs

Plasmids used for production of the minimal-domain NF-Y complex (NF-Ymd) were described in (30). CDSs of mouse PBX1 (aa 1-317), PREP1 (aa 1-344) and Sp2 N-terminus (aa 1-94), including substitution mutants, were codon-optimized and obtained as synthetic gene strands from Eurofins Genomics. The synthetic sequences included homology regions with the expression vectors for NEBuilder assembly reactions. pCoofy1 (6xHis) and pCoofy4 (6xHis-MBP) expression vectors, described in (54), were linearized by PCR amplification using Q5® Hot Start High-Fidelity 2X Master Mix (NEB). Sp2 wt and the designed substitution mutants were cloned into pCoofy4 using NEBuilder HiFi DNA Assembly Master Mix (NEB), resulting in N-terminally tagged 6xHis-MBP fusion proteins. For TALE heterodimer production, PBX1 and PREP1 CDSs, separated by a ribosome binding site, were cloned in this order into pCoofy1 as a bicistronic construct, resulting in the co-expression of 6xHis-PBX1 and PREP1. All constructs were verified by Sanger sequencing.

### Production and purification of recombinant proteins

NF-Ymd heterotrimer expression and purification was described in (30). 6xHis-PBX1/PREP1 heterodimer and 6xHis-MBP-Sp2 constructs were transformed in BL21(DE3) *E. coli* cells. The overnight-grown preculture was diluted 1:50 in LB medium, grown at 37°C and induced with 0.3 mM IPTG at OD_600_ = 1. Cells were harvested 1.5 hours after induction. Induced cell pellets were resuspended in Buffer A10 (10 mM Tris-HCl pH 8.0, 400 mM NaCl, 2 mM MgCl_2_, 10% glycerol, 10 mM imidazole) including protease inhibitor cocktail (GeneSpin). Cells were lysed by sonication and the soluble fraction was incubated with HIS-Select® Nickel Affinity Gel (Sigma-Aldrich) for 1 hour at 4°C. Proteins were eluted in Buffer A containing 250 mM imidazole and dialyzed in Buffer B (10 mM Tris-HCl pH 8.0, 400 mM NaCl, 10% glycerol, 2 mM DTT) using Pur-A-Lyzer Maxi 6000 dialysis kit (Sigma-Aldrich).

### Electrophoretic Mobility Shift Assays (EMSAs)

EMSAs were performed as described in (55). Briefly, reactions were assembled using a 5′ Cy5-labeled dsDNA probe matching the sequence used for AF modelling (**Supplementary Table 1**). Each recombinant protein was added to a reaction binding mix with the following final composition: 20 nM Cy5-labeled DNA, 12 mM Tris-HCl pH 8.0, 50 mM NaCl, 50 mM KCl, 0.5 mM EDTA, 5 mM MgCl_2_, 2.5 mM DTT, 0.1 mg/mL BSA, 12% glycerol. After assembly, reactions were equilibrated at 25°C for 30 min. In direct competition EMSAs, an excess of unlabelled DNA competitor was included in the reaction mix prior to protein addition. Electrophoresis was performed using 4.5% non-denaturing polyacrylamide gels in 0.25× TBE. The gel was run in cold 0.25× TBE at 100 V. Fluorescent gel images were acquired using a Chemidoc MP imaging apparatus (Bio-Rad).

## RESULTS

### Genomic association of PREP1/PBX1, NF-Y and Sp2

It was previously shown that the mouse Sp2 genomic locations overlap the DECA-CCAAT composite motif-GAnnGAC-11bp-CCAAT-occupied by PREP1/PBX1 and NF-Y (**Fig. 1A**). PREP/PBX, NF-Y and Sp2 ChIP-seq peaks are available from human K562 ENCODE experiments, organized by the Factorbook repository (56). The preferred motifs of PBX1 and PREP and of NF-Y subunits are the expected DECA and CCAAT motifs, respectively (**Fig. S1**; see also (24)): the DECA-CCAAT composite motif is found significantly enriched in Sp2 locations (**Fig. 1A**). We decided to focus on the promoter of the mouse *Sp2* gene, itself a validated target of Sp2 (28). Indeed, available ChIP-seq data on mouse embryonic fibroblasts show peaks of all six proteins on the *Sp2* promoter (**Fig. 1B**). The *Sp2* promoter harbours five CCAAT within the −150/+1 region: two are separated from DECA motifs by the canonical distance of 11 bp, a third by 10 bp. The conservation of these elements in mammals is almost absolute (**Fig. 1C**). These data confirm mouse and zebrafish genomic data in human cells (22, 23). We selected the DECA-CCAAT composite motif closest to the TSS of the *Sp2* promoter (−87/–38 region) for further structural analyses.

### AlphaFold 3 model of PREP1/PBX1, NF-Y and Sp2 on the DECA-CCAAT site

We fed AlphaFold 3 with a fragment of 50 bp encompassing the DECA-CCAAT composite motif of the mouse *Sp2* promoter (**Fig. S2**) and the six mouse protein sequences outlined in **Fig. 2A**. Our first attempt to model the entire complex with full-length proteins returned a result that was not reliable, due to evident hallucinations involving the disordered regions of the polypeptides (spurious structural order in the form of α-helices predicted with very low confidence) (**Fig. S3**). Thus, we limited the analysis to amino acids (aa) 1-94 for Sp2, the region previously identified as necessary for chromatin binding (28); for the TALE TF subunits PBX1 and PREP1 we included the heterodimerization domains and DNA-binding HD regions: aa 40-322 for the former, aa 40-320 for the latter (**Fig. 2A**). For NF-Y we used the conserved domains portions involved in trimer formation and CCAAT-binding as in the 4AWL crystal structure: aa 262-332 of NF-YA (HAP2 region, named after the yeast protein), 51-143 of NF-YB and 27-120 of NF-YC (**Fig. 2A**). We did not consider the large, presumably unstructured, QVIT-rich TADs of NF-YA, NF-YC and Sp2.

The AF3 structural model of the complex obtained with the six polypeptides and the *Sp2* promoter DNA is shown in **Fig. 2B**, depicted in colours reflecting the relative pLDDT confidence level (dark blue and light blue being very high and high, respectively, orange being the lowest). In **Fig. 2C**, we outline the individual protein chains and DNA boxes with different colours. The TFs are correctly modelled as heteromeric complexes, associated to DNA through their DBD regions, predicted with high confidence scores (**Fig. 2B**). Instead, Sp2 is tethered to the complex through direct protein-protein interactions with the TALE heterodimer and NF-Y (see below). NF-Y and TALE TFs are composed of mostly helical domains. The DECA motif is bound by the IZ-helices of the two TALE HDs. CCAAT is recognized by the NF-YA DBD, which inserts into the minor groove; additional non sequence-specific contacts are made by the NF-YB/NF-YC HFD dimer, inducing a pronounced bend in the DNA curvature, as expected from NF-Y experimental structures (**Fig. 2C**).

The model predicts with high confidence the topology of the large PBX1/PREP1 heterodimerization module, which was previously unknown. Connected to the HD through linkers of different length in the two subunits, the TALE module adopts a composite and apparently rigid fold, composed of an elongated helix bundle with a globular head (see below), leaning towards the side of the DNA molecule occupied by NF-Y (**Fig. 2C-D**). The long linker region of PREP1 between MEINOX and HD, predicted to be unstructured, is rendered as an extended loop, which protrudes from the otherwise compact heterodimeric TALE structure (**Fig. 2D**).

A general scheme of the reciprocal interaction surfaces within the predicted complex is shown in **Fig. 2E**. Interestingly, Sp2 has two major anchor points with the complex (shown in red): #1 with specific segments of PREP1 and PBX1, #2 exclusively with the N-terminal part of the HAP2 region of NF-YA. These Sp2 anchor points are separated by an unstructured linker region that physically connects the TALE heterodimer to NF-Y (see also **Fig. 2C-D**).

The extended connections between the NF-YB/NF-YC heterodimer and NF-YA are expected, as those between the TALE TFs. The reliability of the global 3D arrangement of the complex (**Fig. 2F**) shows how confidently each region has been spatially positioned relative to other domains/elements within the predicted structure (PAE score): as expected, it is high for NF-Y subunits, but also for PBX1 and PREP1, except for the linker region of the latter. The scores are lower for Sp2, yet the discrete anchor points #1 and #2 are confidently positioned.

In summary, AlphaFold 3 predicts with reasonable confidence numerous details of the DNA-binding TFs in association with Sp2, with the latter predicted to connect TALE to NF-Y. The novel aspects will be thereafter expanded.

### DNA binding of NF-Y to CCAAT and PBX1/PREP1 to DECA

First, we checked the predicted DNA-binding contacts of NF-Y, as these are structurally known. **Fig. S4A** shows an overlay of the mammalian trimer in the AF3 model (dark green) and the crystal structure (grey, PDB: 4AWL), which are essentially superimposable, with limited deviations along their main chain structures. An important feature of the complex is DNA bending: NF-Y does impose a severe curvature of the double helix trajectory in the AF3 model, which is centred on the CCAAT box. 5’ of the CCAAT (left side of the Figure), where the DECA site is located, the DNA axis is not distorted. The nucleobase contacts, essentially established by NF-YA, are shown in **Fig. S4B**: basic residues residing in helix A2 and adjacent loop-R274, H277, R281, R283-make the same contacts with the minor grove of CCAA (nt numbers 29-32), similarly to G286, G287, R288, as well as F289 of the GXGGRF loop region. The AF3 model lacks a couple of contacts from NF-YB A60 and NF-YA R274 on the bottom strand, the former to A<u>T</u>TGG, the latter to nucleotide number 34 (a C in the *Sp2* site but a G in the HSP70 site of 4AWL). In summary, AlphaFold 3 faithfully predicts NF-Y contacts, as well as distortion of the DNA helix around the CCAAT box nucleotides.

We then considered the PREP1/PBX1 homeodomains bound to the *Sp2* DECA motif (GAgcGAC): **Fig. 3A** shows the overlay of PBX1 protomer HD, as observed within the combined AF model, with the available crystal structure obtained with the HOXA9 partner (PDB: 1PUF) in the left Panel; in the right Panel, the corresponding HD of PREP1 is overlayed on the paralog MEIS1 HD homodimeric crystal structure (PDB: 4XRM). Overall, the HD folds in the model are superimposable to the experimental structures. We compared the HD/DNA contacts in the AF3 model (**Fig. 3B**, central Panel) to the respective half-sites of the crystal structures (**Fig. 3B**, left Panel for PBX1 and right Panel for MEIS1). In general, PBX1 contacts are on the left half-site of the motif, recognizing the cGA trinucleotide (nt number 10-12), PREP1 on the right half-site, GACgg (nt number 15-19). The intervening nucleotides, GC in the DECA site used here (positions 13-14), are not contacted and indeed show variability in the DECA sequence logo (**Fig. 1A**). **Fig. S5** shows the organization of PBX1 and PREP1 HDs. The PBX1 contacts are exerted by helix a3 residues N286 and R290, with the N-terminal arm R237 predicted to make side chains interactions with low confidence in the minor groove: the same amino acids are involved in contacts in the crystal structure, with the addition of N282 contacting a T in 1PUF, but missing contact with the G (nt number 13) in the *Sp2* promoter. On the PREP1 side, sequence-specificity is provided by the a3 residues N312, R316 (top strand) and R315 (bottom strand) on the core GAC trinucleotide (positions 15-17), with I311 contacting the C in the bottom strand (position 19). These residues correspond to the conserved I324, N235, R328, R329 of MEIS1 in the dimeric 4XRM crystal structure. The AF3 model predicts-with low confidence-additional minor groove GA contacts of R263 (N-terminal arm of the HD): the MEIS1 corresponding residue-R276-was not included in the construct used in 4XRM. Overall, HD a3 residues known to provide sequence-specificity-N51 and R55 in the homeodomain numbering-are indeed positioned to contact the corresponding bases through H-bonds, both in PBX1 (N286, R290) and PREP1 (N312, R316). Moreover, PBX1/PREP1 DNA-binding was experimentally validated with mutants for the above amino acid residues, including I311 for PREP1, showing significantly impaired DNA association (**Fig. 3B**) (57). We conclude that the AF3 model faithfully places PBX1 and PREP1 HDs on the respective sides of the DECA motif, contacting selected DNA bases with residues previously shown implicated in sequence-specificity in experimental studies of isolated proteins’ domains in complex with DNA. In summary, the AF3 model is credible both on the NF-Y and the PBX1/PREP1 sides and correctly orients the two TFs on DNA.

**Figure 3.**
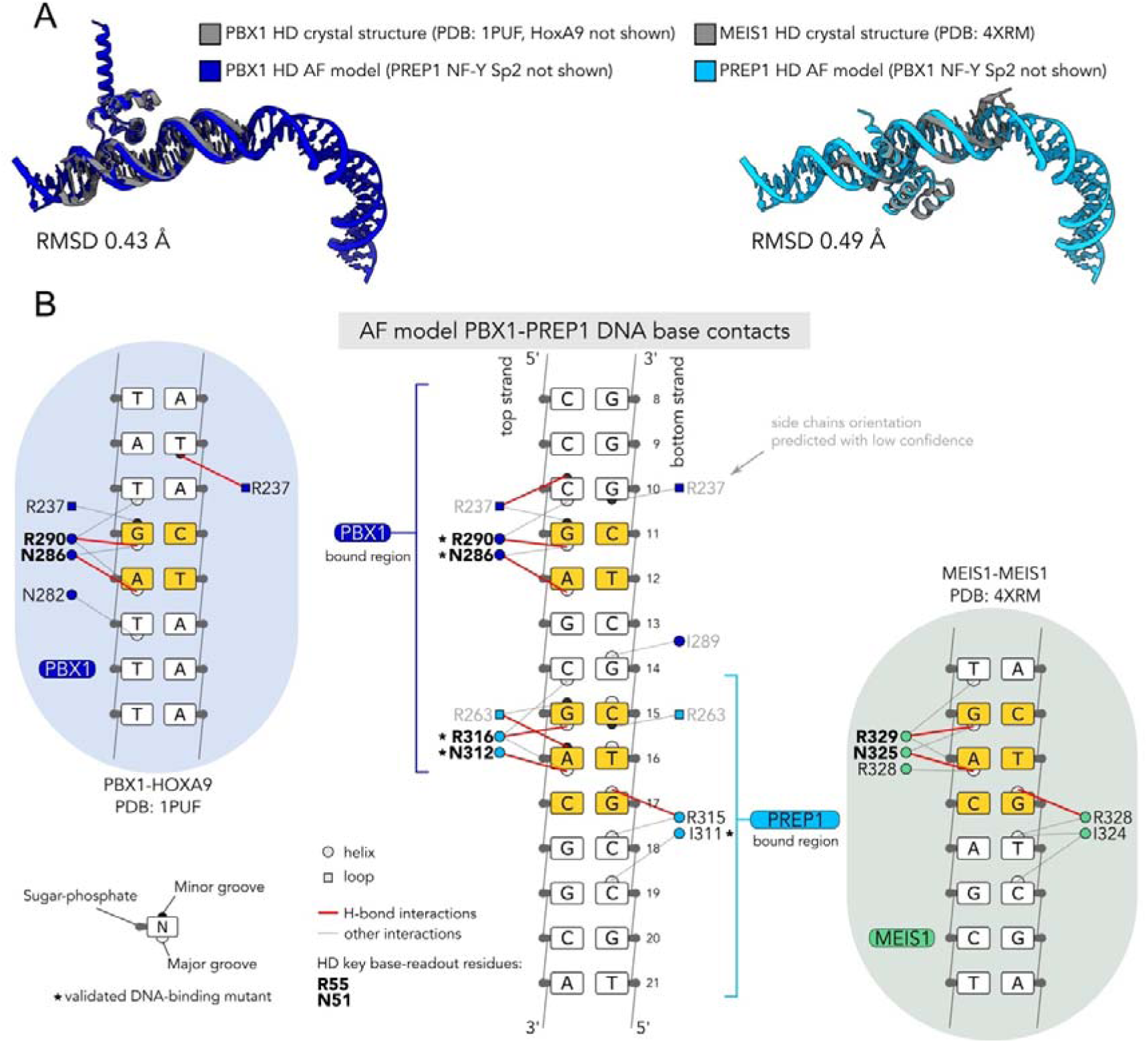
Predicted DNA contacts of PBX1/PREP1 with the DECA motif. **A**. Superimposition of PBX1 HD within the predicted TALE/NF-Y/Sp2/DNA complex on PBX1/HOXA9/DNA crystal structure (PDB: 1PUF, left panel) and superimposition of PREP1 HD with MEIS1/MEIS1 crystal structure (PDB: 4XRM, right panel). Only HDs for each protein and DNA are shown. **B**. Comparison between PBX1/PREP1 predicted contacts with DNA nucleobases in the TALE/NF-Y/Sp2/DNA complex model (central scheme) and the available crystal structures shown in A (left and right inserts). For schemes referred to crystal structures, only the relevant protomer recognizing the correspondent DECA half-site is shown. DNA bases positions constituting the two DECA half-sites are highlighted in yellow. Direct contacts with DNA nucleobases are indicated by red (H-bonds) or dashed lines. Contacts with DNA sugar-phosphate backbone are omitted for clarity. Residues with validated DNA-binding mutants are indicated by asterisks. Protein residues contacts predicted with low confidence scores are labelled in light grey colour for the AF model.

### The PBC-MEINOX domains form the predicted all-helical heterodimerization module

The heterodimerization interfaces of PBX1 and PREP1 predicted by AF3 overlap with the corresponding PBC and MEINOX domains (**Fig. 2**), known to be necessary and sufficient for complex formation. The model displays a composite topology, displaying an elongated architecture constituted by five confidently predicted IZ-helices of PBX1 (H1-5) and four of PREP1 (H1-4) (**Fig. S6**), generally organized in two subdomains: an elongated ‘stem’ region, composed by an intermolecular coiled-coil core formed by three helices, joined to a globular ‘head’ formed by a bundle of helical elements associating the N-terminal portion of the long PREP1 helix H4 (**Fig. 4A**). The head region is constituted of the N-terminal helix H1 and 2 of PBX1, as well as helix H1, 2 and 3 of PREP1, together with the N-terminal end of H4.

**Figure 4.**
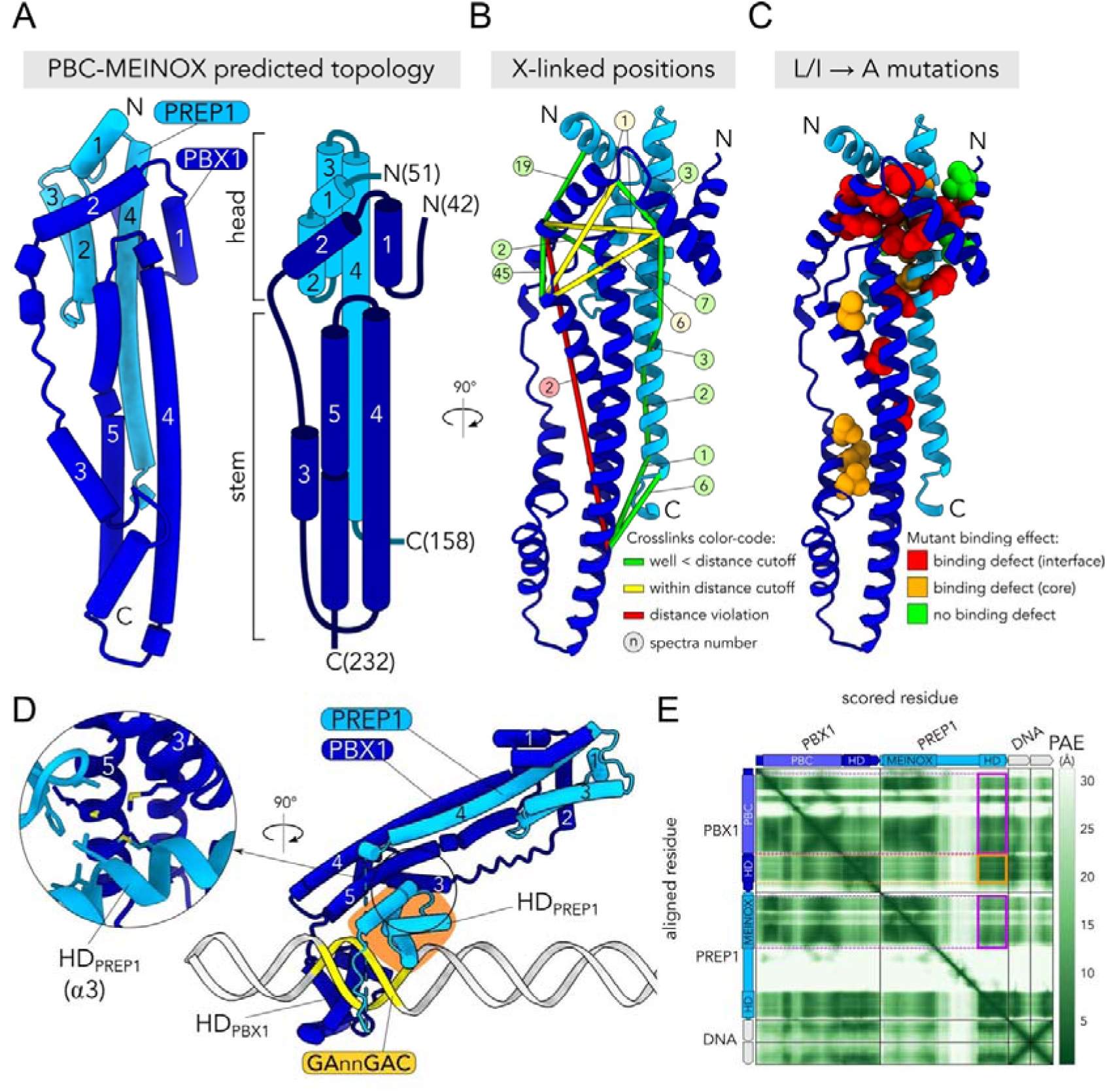
Structural features of the predicted PBX1-PREP1 heterodimerization module (PBC-MEINOX domains). **A**. The overall topology of the PBX1-PREP1 heterodimerization module in the predicted model (left) and a simplified cartoon (right). Each main helix is numbered from N-to C-terminal position. **B**. Cross-validation of the predicted PBC-MEINOX domain structure with experimental data obtained with in-solution XLMS on the purified complex. Ca of cross-linked Lys residues are connected by lines coloured based on distance cut-offs (green and yellow for residues below the compatible distance cut-off). The number of XLMS spectra detected for each residue pair is shown. **C**. Mapping of experimentally tested mutations for PBX1-PREP1 dimerization on the predicted structural model. Side chains tested with alanine mutants are shown as space-filled residues and coloured according to the mutant effect on dimerization and whether they lie at an inter-(red) or intra-(orange) protein interface. Cross-linking and mutant data are from (13). Protein sequences used for modelling match those used for cross-linking experiments (PBX1 aa 1-308; PREP1 aa 1-344). Only the PBC/MEINOX domains are shown for clarity (PBX1 aa 42-232; PREP1 aa 51-158). **D**. AF3 model of PBX1/PREP1 heterodimer bound to the mouse *Sp2* promoter DECA site. PREP1 HD is highlighted in orange and the interaction surface with PBX1 is expanded in the circular inset. **E**. PAE heatmap of the model shown in D. Violet rectangles highlight that the PBC-MEINOX domains are confidently positioned relative to PREP1 HD.

Specifically, PREP1 helix H2, 3 and 4 form a short three-helix bundle, which together with PREP1 helix H1 present a hydrophobic surface onto which helix H1 and H2 of PBX1 are docked. The main scaffold of the stem region is composed of the two contiguous amphipathic helices H4 and 5 of PBX1 folded in a long α-helical hairpin providing a hydrophobic docking site for the C-terminal end of PREP1 helix H4. The short PBX1 helix H3 interacts with the opposite side of the α-helical hairpin. The head and stem regions are rigidly connected through the long PREP1 helix H4 spanning the two subdomains. Finally, PBX1 helix H1 and 2 in the head region interact with the loop separating PBX1 helix H4 and 5 in the stem. The axis of PBX1 helix H5 deviates around S210, avoiding clashes with PREP1 helix H2. In summary, the predicted PBC-MEINOX heterodimerization module shows a rigid and elongated core structure constituted by the head and stem regions, flanked by the poorly structured hydrophilic linker connecting PBX1 helix H2 and 3. We retrieved no significant matches for similar protein folds in experimental structures present in the PDB database using Foldseek (58), suggesting that TALE heterodimerization module may represent a novel heterodimeric assembly mode with no known structural homologies.

We verified the robustness of the AF predicted structure of the PBC-MEINOX domains heterodimeric fold by examining published data from both cross-linking mass-spectrometry (XLMS) and site-specific mutagenesis experiments obtained with purified PBX1/PREP1 in solution, in the absence of DNA (13).

We first mapped XLMS positions on an AF model we built using protein sequences matching those used in the experiment (**Fig. 4B** and **Supplementary Table 3**). Overall, 12 of the 13 crosslinked positions within the PBC-MEINOX domains are below the crosslinking distance cut-off, suggesting adherence of the structural module with the experimental data. On the other hand, crosslinks involving regions outside the PBC-MEINOX domains (not shown) are generally above the distance cut-off in the model. This is likely caused by the flexibility of the HDs and linker regions relative to the PBC-MEINOX module in absence of DNA (as in the experiment) and by the limit of AF to return a number of sufficiently representative interdomain conformations.

We then mapped the positions of PBX1 and PREP1 mutated in heterodimerization tests performed with recombinant proteins (13): we found that 23 of the 24 residues proven to have significant binding defects when mutated are either located at the PBX1/PREP1 interface, impacting on affinity directly (15/23), or at the core of intrasubunit contacts (8/23), thus contributing to the proper folding of the protein (**Fig. 4C** and **Supplementary Table 4**). In addition, the positions of seven mutants showing no binding defects are either solvent exposed (1/7) or lie in disordered/low-confidence regions (6/7). We conclude that the intersection with experimental data supports the structure of the TALE heterodimerization module predicted by AF.

In the model bound to DNA, the PBC-MEINOX heterodimerization module leans on top of the PREP1 HD (**Fig. 4D**, see also **Fig. 2C**). The interaction is mediated by helices H3 and 5 of PBX1 PBC domain, contacting PREP1 HD a1 and a3. The relative interdomain distances are predicted with high confidence (**Fig. 4E**). This results in the evident inclination of the PBC-MEINOX module towards the DECA DNA side occupied by PREP1. Flipping the half-sites within DECA sequence on opposite strands indeed reorients the TALE heterodimer (**Fig. S7**), highlighting the importance of sub-motifs’ orientation within the DECA-CCAAT composite sites.

In conclusion, PBX1 and PREP1 heterodimerize using a complex structural module with an architecture that might be unique to the TALE family. The model also predicts that this module is the docking site for the SP-box motif in Sp2, as described next.

### Sp2 bridges PREP1/PBX1 and NF-Y on the canonical DECA-CCAAT composite motif

There are several aspects of the specific interactions of Sp2 with PBX1/PREP1 and NF-Y worth detailing (**Fig. 5A**). Two Sp2 subdomains show a high pLDDT score working as anchor points for association with the DNA-bound TFs (see also **Fig. 2**). The first subdomain corresponds to the SP-box (aa 34-45), shared by the other Sp family members. It forms an α-helix binding the PBC-MEINOX heterodimerization module stem region, running parallel over PREP1 helix H4 and antiparallel to PBX1 helix H4, extensively contacting both in a surface groove (**Fig. 5B**, left Panel). Although some H bonds are predicted-E174 of PBX1 with T41 and K44 of Sp2-most of the contacts are hydrophobic (**Fig. 5B** and **Fig. 5D**, right Panels). The second subdomain is related to contacts with NF-Y, which are more limited, involving a short stretch of Sp2 (aa 68-73 **Fig. 5C**). The contacts involve exclusively a hydrophobic pocket formed by the most N-terminal portion of NF-YA conserved core. At this site, Sp2 is predicted to form a 3 aa-long two-stranded parallel β-sheet with NF-YA aa 264-266, sustained by main chain H-bonding, namely involving Sp2 P69-K71-A73 and NF-YA L264-V266 (**Fig. 5D**, left Panel). We name this novel Sp2 region YA-SLiM (for NF-<u>YA</u> binding <u>S</u>hort <u>Li</u>near <u>M</u>otif) to indicate the NF-YA docking capacity. Modelling of the Sp2 1-94 region anchor points separately with the two partner TFs using AlphaFold2-multimer returns the same interaction sites with very high confidence (**Fig. S8**). Both Sp2 stretches are perfectly conserved across mammals, amphibians and fishes (**Fig. 5E**). The SP-box and YA-SLiM motifs are embedded in a protein region predicted to be intrinsically disordered, occupying distinct evolutionarily constrained regions (**Fig. S9**). Yet, YA-SLiM is not present in other Sp-family members (**Fig. S10A**): indeed, substitution of Sp2 with Sp1-lacking YA-SLiM-in AF3 modelling leads to loss of anchor point #2, while the interaction with TALE is maintained through the conserved SP-box (**Fig. S10B**). In the same model, we noticed that the disordered PREP1 linker region occupies the NF-YA pocket otherwise occupied by Sp2, although at very low confidence and with no secondary structure elements, potentially representing a spurious modelling result. The stretch than links the SP-box to YA-SLiM in Sp2 proteins (**Fig. 5E**), visibly less conserved in amino acid sequence but not in length (20 residues), scores low in structure prediction. It is imaginable that the linker is deemed to be intrinsically flexible to accommodate potentially different distances between PBX1/PREP1 and NF-Y. Overall, the AF3 models are compatible with previous biochemical studies regarding the role of Sp2 N-terminal portion in interacting with, and stabilizing, the TALE/NF-Y complex on DECA-CCAAT composite motifs (28).

**Figure 5.**
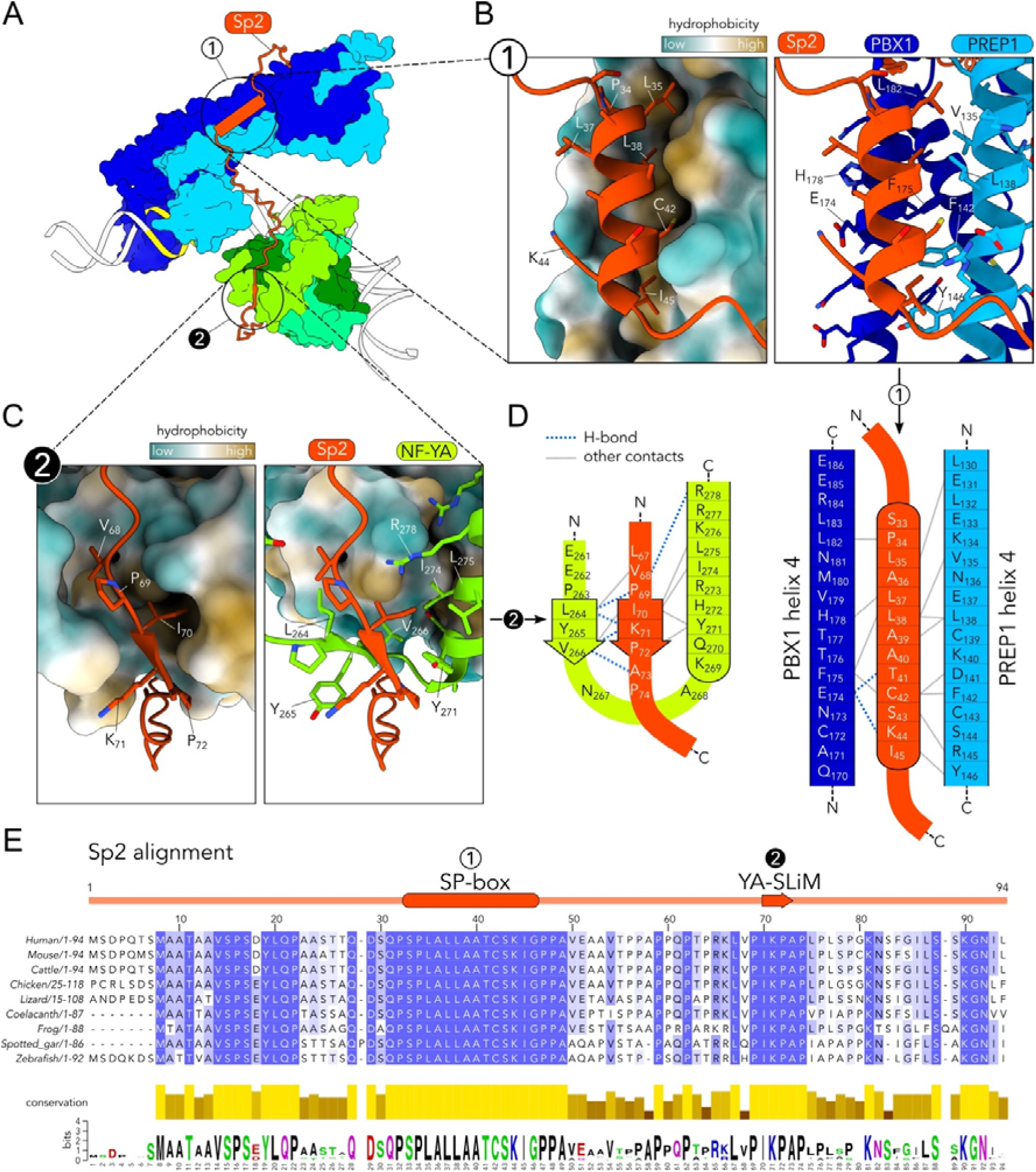
Sp2 anchor points on PBX1/PREP1 and NF-Y. **A**. Overview of the predicted Sp2 anchor points on the PBX1-PREP1 heterodimerization module (#1) and on NF-Y (#2). **B**. Interaction surface of anchor point #1 constituted by the SP-box with a hydrophobic groove composed by the PBC/MEINOX domains. Residues predicted to be involved in the interaction are indicated. **C**. Same representation as in B, but for anchor point #2, constituted by a short linear motif of Sp2 interacting with a hydrophobic pocket formed by NF-YA. **D**. Schematic representation of the predicted interactions of anchor points #1 and #2 in the model. **E**. Multi-species sequence alignment of Sp2 N-terminal region (1–94) used in modelling. Resulting conservation scores and sequence logo are shown below the alignment.

### Reconstruction of the TALE/NF-Y/Sp2/DECA-CCAAT complex with recombinant proteins

To validate the structural predictions, we reconstituted the complex bound to DNA *in vitro*. For NF-Y and Sp2 we used the domains matching those of AF3 modelling. For the TALE heterodimer we used regions reported to be amenable for protein purification (PBX1 1-317; PREP1 1-344) (59). The recombinant proteins were produced in *E. coli* and purified (**Fig. S11A**). The TFs were tested in EMSAs using a Cy5-labelled oligonucleotide corresponding to the *Sp2* promoter DECA-CCAAT DNA used for predictions. NF-Y and TALE form distinct and well-defined low-mobility DNA complexes in the gel (**Fig. S11B**). The combination of the two TFs results in the formation of a ternary complex with lower mobility, which seems to form more readily than the binary complexes, suggesting that binding to DNA is favoured by the co-presence of NF-Y and TALE (**Fig. S11B**). Addition of Sp2 increases NF-Y/TALE ternary complex abundance and subtly shifts its migration in the gel, suggesting the formation of the quaternary complex. Instead, Sp2 is incapable of generating a DNA complex alone, or individually with NF-Y or TALE (**Fig. S11B**). These complexes are competed efficiently by an unlabelled DECA oligo, proving that DNA-binding is specific (see below). To verify the relevance of SP-box and YA-SLiM in formation of a stable complex, we designed Sp2 mutant proteins containing single amino acid substitutions to a Gly residue either in the SP-box (mL; L38G) or YA-SLiM (mI; I70G), as well as the double mutant in both motifs (mLI; L38G, I70G) (**Fig. 6A**). The targeted residues correspond to direct hydrophobic contacts predicted in the structural models (see **Fig. 5B, C**). We challenged wt and mutant Sp2 to form a complex with the NF-Y/TALE/DNA assembly (**Fig. 6B**): formation is observed with the wt Sp2, at the lowest dose (compare Lanes 4 with 5-7), but not with mLI nor mL mutants (Lanes 8-10 and 14-16, respectively). Addition of mI, the Sp2 mutated in YA-SLiM, does yield a slower complex, but only at the highest dose (Lane 13). The residual binding of this mutant might depend on the backbone-driven H-bonding that characterize the YA-SLiM interaction spot. To substantiate the properties of the NF-Y/TALE/Sp2 complex, we performed competition experiments with a cold DECA oligo (**Fig. 6C**). As expected, NF-Y was not competed (Lanes 2 and 3), whereas TALE and NF-Y/TALE complexes were readily competed (Lanes 4-7). Instead, the slower-mobility complex generated by addition of wt Sp2 polypeptide was minimally competed, unlike the faster NF-Y/TALE complex in the same reaction, which was abolished (Compare Lanes 8 and 9). Addition of the three Sp2 mutants did not modify the mobility of NF-Y/TALE bands which were efficiently competed (Lanes 10-15). This is a further proof of stabilization of the NF-Y/TALE complex on DNA by the wt, but not the mutated, Sp2 protein.

**Figure 6.**
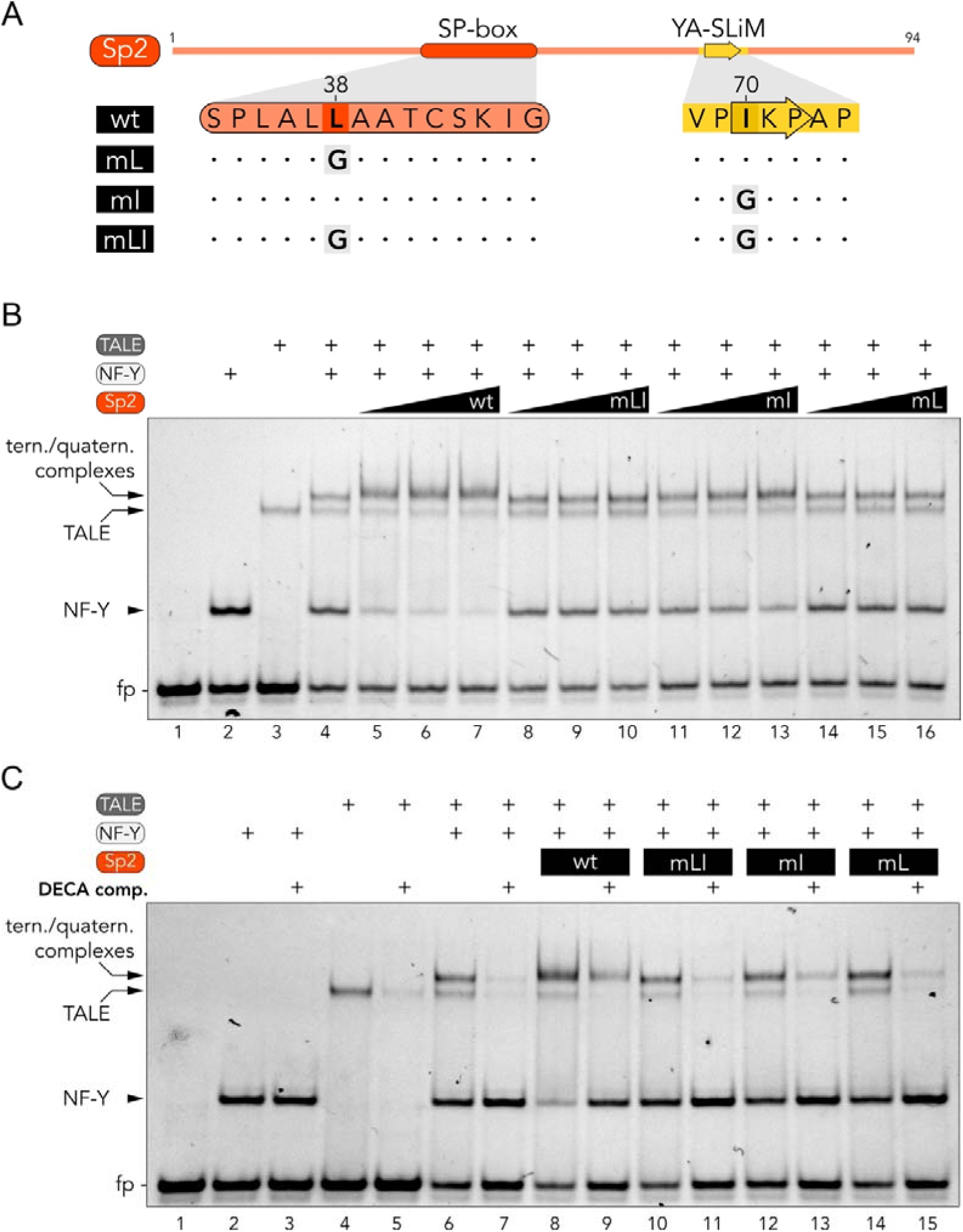
*In vitro* validation of Sp2 anchor points assessed with EMSA experiments. **A**. Schematic representation of Sp2 single amino acid substitution mutants to Gly in SP-box (mL), YA-SLiM (mI) or both (mLI) motifs. **B**. Sp2 dose-response EMSA experiment. Protein concentrations in the assays are the following: 40 nM NF-Y, 60 nM TALE and increasing amounts of Sp2 proteins (160, 320, 640 nM). DNA probe concentration is 20 nM. The different DNA-bound complexes and the free probe (fp) are indicated on the left. **C**. Competition EMSA experiment. Reactions were assembled in the presence of a 10-fold excess of unlabelled DECA competitor DNA (DECA comp.) or TE as control. Same protein concentrations as in B (320 nM Sp2).

From these DNA-binding assays with recombinant proteins, we conclude that (i) binding of the TALE TFs to the DECA site is facilitated by NF-Y; (ii) Sp2 associates to the ternary complex, requiring the intact SP-box and YA-SLiM subdomains; (iii) this interaction leads to the cooperative stabilization of the NF-Y/TALE complex on DNA. These experimental observations validate the role of Sp2 as suggested by the AF3 models described above.

### Prediction of DNA rules for the association of the TALE/NF-Y/Sp2 hexameric complex

Having modelled and confirmed *in vitro* the multiprotein/DNA complex, we tested it by varying the natural distance of 11 bp between DECA and CCAAT on the *Sp2* promoter DNA (**Fig. 7**, central Panels). In the right Panels we show the average pLDDT confidence score for Sp2 at each distance, compared to the canonical 11 bp spacer. By increasing it of 1 bp (**Fig. 7**, 12 bp spacing, lower central Panels), the Sp2 linker is predicted to adopt a more extended conformation (side views of 11 *vs* 12 bp), adapting to the increased distance between the DNA-bound protein complexes, as the SP-box and YA-SLiM anchor points remain in place and similarly well-structured as in the wt DNA (Lower right Panels). Instead, addition of 2 bp-distance of 13 bp and consequent rotation by ∼70°-leads to detachment of anchor point #2 from NF-Y, with YA-SLiM modelled as fully unstructured (see pLDDT plot, right Panel), suggesting that the b fold arrangement is adopted only upon NF-YA association. The SP-box IZ-helix appears instead to be correctly positioned and structured on PBX1/PREP1 (**Fig. 7**, lower Panels), potentially because of the more extensive interactions compared to those of YA-SLiM. Reduction of the DECA-CCAAT distance by 1 bp to the 10 bp configuration results in a ∼30° rotation of the TALE complex along the DNA axis towards NF-Y (**Fig. 7**, upper Panels). In the model, this rotation is accompanied by a slight reduction in DNA bending angle induced by NF-Y, possibly due to the consequent steric hindrance between protein domains. Without this slight DNA rearrangement, the HD of PREP1 would clash with NF-Y, in correspondence of NF-YC a3 helix. Nevertheless, Sp2 is predicted to remain anchored to the complex, with only a slight reduction in the pLDDT score for the YA-SLiM. It is possible that in this configuration the DNA is under some torsional stress to avoid the steric hindrance of the two TFs. In the 9 bp spacing configuration, AF3 still positions TALE and NF-Y on their respective sites, yet the DNA bending induced by NF-Y is further reduced to avoid a full clash of the DBDs of the two TFs. Sp2 maintains its connections, but with a consistent drop in confidence for the YA-SLiM region. An 8 bp distance is fully incompatible with complex formation because of TFs clashing: AF3 predicts the PBX1/PREP1 heterodimer position as not binding to DECA motif, unlike NF-Y which maintains binding to CCAAT. On the other hand, the SP-box α-helix is still predicted to contact PBX1/PREP1, although the overall model scores drop consistently (**Fig. 7**, top Panels).

**Figure 7.**
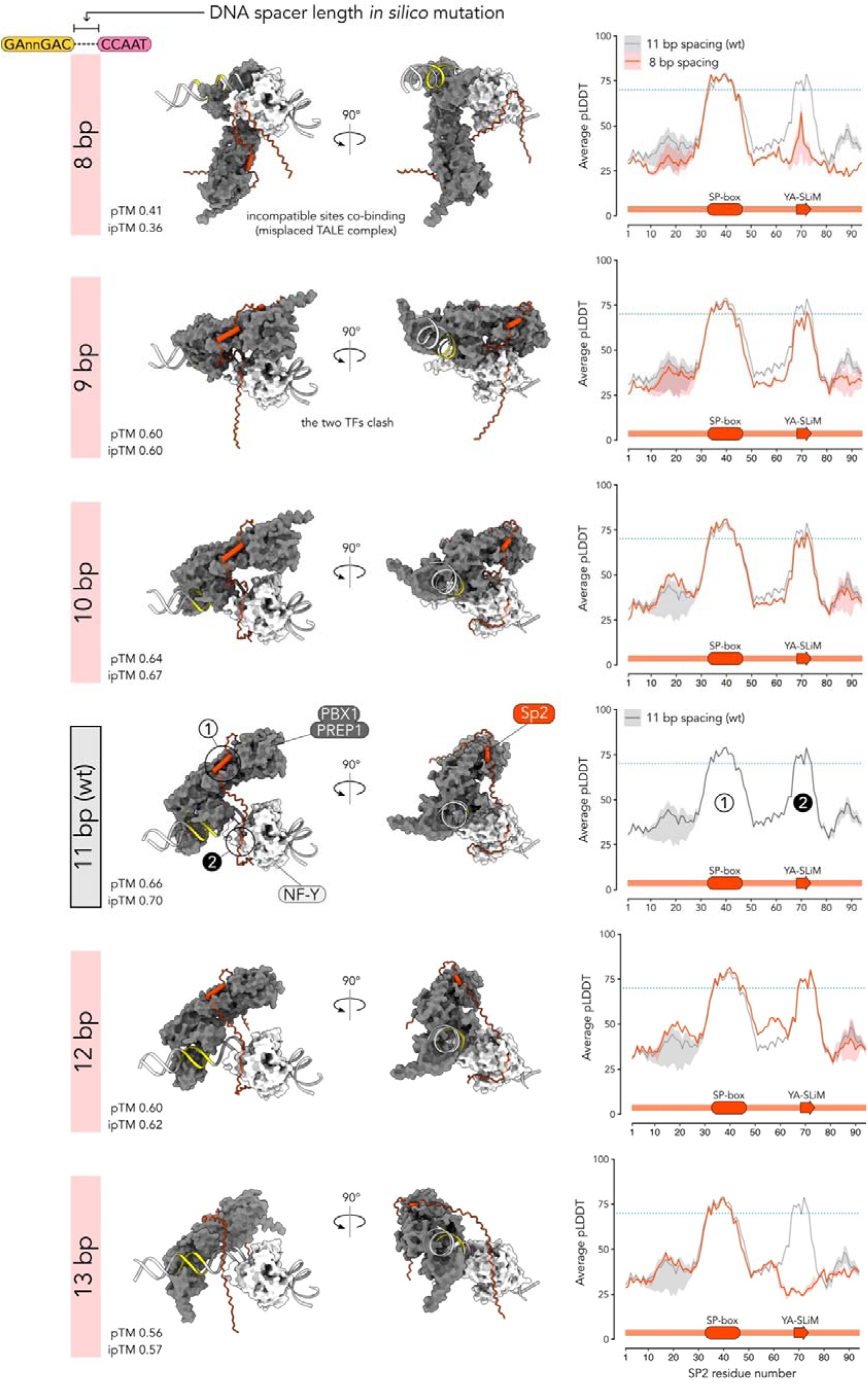
Models with different DNA spacer lengths between DECA and CCAAT motifs. (Left Panels) Frontal and side views of the AF models for TALE/NF-Y/Sp2/DNA complexes with different DECA-CCAAT motifs spacing (the canonical 11 bp spacing model is shown as reference). The surface models of TALE (PBX1/PREP1) and NF-Y complexes are shown in dark grey and white, respectively. Sp2 (aa 1-94) is shown in orange. The unstructured linker of PREP1 is not shown for clarity. The pTM and ipTM quality scores are shown for each model. (Right Panels) Plots of the pLDDT confidence score of Sp2 from the five AF models from each prediction are shown for each DNA spacing configuration (line = mean; shaded area = range). The canonical 11 bp (wt) plot is overlaid as reference (in grey). A dotted light blue line corresponds to pLDDT = 70.

We conclude that the AF3 model likely reproduces the structural basis for the 11 bp optimal distance requirement between DECA and CCAAT recorded by genomic analyses. A reduction in DNA spacing is predicted to make the two TFs clash (severely on the 9 and 8 bp spacings). Instead, an increase in spacing beyond 12 bp is predicted to allow independent DNA-binding of the two TFs, but not simultaneous Sp2 association, thus abrogating its cooperative effect.

## DISCUSSION

We report here on the visualization *via* AF of a multi-protein/DNA complex based on previous genetic characterization of stereo-aligned DECA-CCAAT elements and the biochemical features -and activities-of TALE, Sp2 and NF-Y, as further validated here with recombinant proteins. There are several advancements, expanding current models on how TFs might synergize on DNA, and on our knowledge of individual domains of the TFs considered.

### Geometry of sites and cooperative DNA-binding

Promoters -and enhancers-contain multiple sites for TFs, whose precise spacing is often crucial for function. This means that “appropriate” TFs, among members of often large families of activators (or repressors), need to find a way to coexist in a constrained space. Several models have been so far described. A first 3D paradigm of multiple TFs bound to stereo-aligned sites was provided by the β-IFN promoter, identified by genetic experiments and bound by synergizing NFkB (RelA/p50), IRF3/IRF7, ATF2/JUN (4). Crystallization, modelling and extensive mutagenesis of the enhanceosome super-complex did not reveal crucial protein-protein contacts. At the DNA-binding level, an important stabilizing mechanism is exerted by induction of-subtle-DNA bending by IRF3 to increase ATF2/JUN affinity (i.e. cooperativity mediated by changes in DNA shape). Co-function of the TFs would be provided by TADs mediating recruitment of p300/CBP coactivators, non-DNA-binding platforms integrating contacts with multiple DNA-bound TFs (60). Crystallographic studies provided other examples, which are substantially different. One is NFAT in complex with FOS/JUN, binding to partially overlapping sites of the IL2 promoter: the TFs do make protein-protein contacts crucial for stabilization of the complex, through the Leucine-Zipper of FOS/JUN and the RHR-C domain of NFAT (61). This modality has been recently expanded for additional TFs, indeed using AF modelling (1). Another model was described for ETS family members synergizing with TFs harbouring structurally different DBDs (PAX5, IRF4 and SRF): complexes are stabilized by protein-protein contacts provided by aminoacids involved in DNA-binding: in essence, specific residues can be used to either stabilize binary ETS/DNA contacts, or as docking spots for neighbouring TFs in case of appropriate-often overlapping-geometry of sites (62).

As for NF-Y, several TFs intersect CCAAT with a precise alignment of sites, such as in the E-box–CCAAT configuration (24). We showed that the bHLH USF1/2 and MAX, but not MYC, co-bind *in vivo* to promoters and repetitive sequences (55, 63–65). Biochemical dissection indicates that direct protein-protein contacts are mediated by the USF-Specific Region (USR) with the DBDs of NF-Y. MAX binds efficiently next to NF-Y *in vitro*, but no cooperativity in DNA-binding is observed since it lacks the USR. The domain is short, predicted to be intrinsically disordered and sufficiently flexible to accommodate the 10-12 bp difference recorded between the sites in the genome; shorter or longer distance would bring clashes between the DBDs, or incapacity for the USR to stabilize DNA-binding, respectively (55).

The data of **Fig. 6** suggest that binding of TALE to DECA sites is facilitated by the presence of NF-Y, but direct protein-protein contacts are not confidently discerned in the AF3 models. A possibility to be considered is that this is mediated by the large distortion of the DNA generated by binding of NF-Y to CCAAT, according to the “change in DNA shape” model mentioned above.

We provide predictive details of the TALE/NF-Y/Sp2 complex, also derived from genomic locations and functional studies on frequencies of co-bound TFs, and conserved geometry of DECA and CCAAT. The model described represents yet another way of achieving cooperativity through a stabilizing effect provided by a third TF, Sp2, thus different from the examples mentioned above. The model posits that protein-protein contacts between the DNA-bound TALE/NF-Y are absent-on the 11 bp spacing- or not significant-on the 10 bp spacing, while DNA-binding cooperativity originates from the co-association of Sp2. The assumption has the following caveats: first, we cannot exclude that the disordered linker of PREP1, whose rendered extended trajectory is not to be considered reliable by the low pLDDT and PAE confidence scores, could additionally contact the NF-Y trimer; second, interactions between PREP1 and NF-Y were reported by co-IP (22), which might result from contacts of other parts of the TFs or by indirect interactions. In general, we cannot exclude the possibility that domains not analysed here, such as the respective TADs, could further stabilize the complex. Yet, genomic and biochemical data, conservation of Sp2 regions, the confidence levels of the predicted structures and the EMSA experiments provided here indicate that Sp2 is indeed crucial for the stabilization of the complex. What is implicit in this model (which only considers the N-terminal domain) is that the C-terminal ZF DBD of Sp2 would still be available for further (sequence-specific) DNA contacts, either in proximity of the DECA-CCAAT complex or at a distance, potentially with an enhancer located kilobases away.

### Lessons for the PBX1/PREP1 TALE heterodimer

There are two major advancements potentially derived from the AF3 models, concerning the DNA-binding of the heterodimer HDs to DECA and the structures of protein domains mediating heterodimerization.

AF3 predicts that the DECA nucleotides contacted are those of the PBX1/PREP1 preferred binding site-as found *in vitro* and *in vivo-* and the amino acids modelled within the HDs of the two TFs are typical sequence-specific contact points in isolated PBX1, MEIS1, as well as in other HD-containing TFs (66). The respective positioning is PBX1 at the 5’ half-site, and PREP1 at the 3’ half-site of the asymmetric DECA motif, with all core DNA positions contacted in the heterodimer. Importantly, the models also explain the obligate stereo-alignment of the two asymmetric motifs: the positioning of PREP1 on the DECA half-site (GAC) proximal to CCAAT, and the apparently rigid asymmetric arrangement of the complex, orients the TALE heterodimerization module towards NF-Y, thus favouring the simultaneous association of Sp2 with both TFs.

The TALE heterodimerization module modelled by AF in a novel structural fold is fully compatible with the available biochemical data, both in terms of the protein regions involved and in its predicted topology, and this might have potentially relevant impact on cancer biology. Indeed, the PBC surface of PBX that mediates dimerization with PREP1 is predicted to be shared with the related MEIS1, a powerful oncogene, notably in the hematopoietic system (16). Overexpression of MEIS1 drives transformation of lymphocytes and myeloid precursors, and high levels are found in patients with AML and ALL (67). In addition, MEIS1 appears to be a crucial target of MLL fusion proteins derived from rearrangement of the histone methyltransferase MLL1 with AF9, AF4 or ENL in childhood mixed leukaemia (67).

An oncogenic role has also been suggested in some solid tumours. Whenever tested, mutations in the PBX-dimerization domain abolish oncogenic activity: thus, structural knowledge of the interaction surfaces as presented here could potentially drive searches for peptidomimetic drugs or other small compounds that could interfere with the association (9).

Another field that could be impacted is plant biology: several members of KNOX/PBX (BELL/GLX) heterodimers were isolated decades ago (5, 68) and individual genes shown to drive embryogenesis, including the haploid-to-diploid transition and different developmental processes in land plants (69, 70), as well as in algae (71). The present data should help rationalize the conservation of the PBC/MEINOX domains, as genomic identification of family members proceeds in a rapidly growing number of plant species.

### Lessons for the Sp family

Association to CCAAT-promoters *in vivo* and facilitation of PREP1 and PBX1 recruitment is mediated by the Sp2 N-terminal, not the C2H2 ZF DBD (27). *Dictu* of the availability of the DBD for additional DNA contacts in the multiprotein complex, our findings rationalize the chromatin interactions found in genomic experiments with *Sp2* KO mESCs. In particular, our data suggest a function for the so far elusive SP-box. The SP-box was noticed decades ago at the N-terminal of the founding members of the family and absent in the related KLFs (25, 72, 73). Alignment of SP-boxes highlights significant conservation, which can be reasonably extended to include two additional ZF TFs, ZNF503 and ZNF703 (**Fig. S12**): thus, it is possible that the composite PBC-MEINOX protein fold might be the direct molecular target of other SP-box containing proteins.

The second relevant point is the novel YA-SLiM (VPIKPA), mediating NF-YA contacts. This motif is evolutionarily conserved in the Sp2 of vertebrates, but apparently absent in other members of the family. This short sequence, in complex with NF-Y, adopts an extended b conformation establishing mostly hydrophobic interactions and main-chain H-bonds when complexed with NF-YA HAP2 N-terminal region. From the modelling exercise shown in **Fig. 7**, it is presumed that YA-SLiM loses capacity to interact with NF-Y upon extension of the DECA-CCAAT distance beyond 12 bp, unlike SP-box/TALE interactions, predicted to be more extended and stable. This fits with *in vitro* results obtained with recombinant proteins, showing recruitment of the N-terminal part of Sp2 to a PBX1/PREP1/DECA complex even in the absence of NF-Y (28). The *in vitro* data of **Fig. 6** indicate that both SP-box and YA-SLiM are required for association, as suggested by mutagenesis of the individual elements. In addition, the data of **Fig. 7** implicate the linker region between SP-box and YA-SLiM in restricting the possibility to bridge TALE and NF-Y within a relatively strict window of rotational arrangements (i.e. distances of maximum 1 bp from the canonical 11) along DNA. The related Sp1, which lacks this motif, is known to be one of NF-Y’s best companions, with many examples of juxtaposed sites and cooperative activation of promoters (74, 75).

Protein-protein contacts-through the respective QVIT-rich TADs-were reported (76): thus, Sp family members might interact directly with NF-Y though different domains. These findings pose the obvious question as to whether SP-box and YA-SLiM can connect other TFs, unrelated to TALE and NF-Y: the genomic overlap of sites between the three proteins is large (20/40%), but far from absolute, and bridging other TFs appears a possibility.

### Lessons for NF-Y

The AlphaFold 3 predictions of the multiprotein/DNA complex are consistent with what is structurally known for NF-Y, in terms of nucleotides contacted and amino acids involved. This was expected, given the profound knowledge on 3D structures, basically serving as “positive” internal control for the modelling experiments. There is one novelty, however: the short segment of NF-YA that docks Sp2 YA-SLiM. This peptide stretch is a flexible loop located before NF-YA helix A1, not involved in contacts with NF-YB/NF-YC, yet the amino acids are evolutionarily conserved. They are mostly hydrophobic and centred on Tyr265, which is solvent exposed in the crystallographic structures of NF-Y(CBC)/CCAAT. Interestingly, this residue is phosphorylated *in vivo* (PhosphoSitePlus database: https://www.phosphosite.org) and thus interactions could be potentially regulated by-yet unknown-tyrosine kinases.

### Limitations of AlphaFold modelling in studying TF/DNA complexes

Despite the evident advancements provided by AF3 to model protein-DNA complexes, confirmed here by biochemical means with recombinant proteins, several limitations must be taken into account in interpreting the results. (i) Modelling large multiprotein/DNA complexes remains problematic, due to the low reliability of the results (hallucinations) when feeding AF3 with large amount of tokens (>∼2500), especially with proteins rich in intrinsically disordered regions (IDRs) (43). Many of the complexes involved in gene regulation easily exceed this size limit. Our analyses are limited by this factor, since we identified the “minimal” domains that recapitulate the available experimental observations. Yet, we cannot exclude that interactions mediated by regions excluded in our models might play a role in the assembly. (ii) A second issue is given by the absence of explicit biophysical energetic parameters in the construction of AF3 models. The models do not provide a direct way of assessing the strength of a biomolecular interaction. For instance, AF3 tends to position a DNA-binding protein on any DNA molecule (if available) even in the absence of its specific recognition site, although with low confidence levels. While this behaviour can be interpreted as a surrogate for the natural non-specific DNA-binding activity of any DNA-binding protein at high-enough concentrations, it does pose issues when interpreting models for proteins with unknown binding motifs. Additionally, when reconstructing multi-TFs complexes on DNA, the relative position and confidence of each of them should be carefully examined and confronted/confirmed with experimental data. (iii) The disordered regions modelled by AF3 fail to faithfully sample the space of their natural conformational ensembles. The same is true for proteins characterized by dramatic conformational changes, often important for their activity. This limits the interpretation of the overall size/conformation of a specific disordered region in AF3 models. Consequently, it is not clear whether AF3 can predict direct interactions mediated by two IDRs, a scenario likely extremely common among TFs.

### Perspectives

Recent systematic AI-driven genomics predictions identified CCAAT and GC boxes among the few TF binding sites impacting on TSS positioning in coding and non-coding genes (77–79). Although it is not yet possible to model large pre-initiation complexes that include most players required for promoter activation, the present approach of dissecting an NF-Y-based multicomponent DNA assembly could be extended to other TFs known to synergize-Sp1, RFX, CREB, ATF6, HSF1, SREBP, USF1-as well as any other TF combinations, even in cases in which 3D knowledge is still partial.

## AUTHOR CONTRIBUTIONS

Andrea Bernardini: Conceptualization, Investigation, Formal analysis, Methodology, Visualization, Data curation, Validation, Writing—original draft, Writing—review & editing. Nerina Gnesutta: Conceptualization, Visualization, Writing—review & editing. Roberto Mantovani: Conceptualization, Visualization, Writing—original draft.

## SUPPLEMENTARY DATA

Supplementary Data are available in the online version of the article.

## DATA AVAILABILITY

ChIP-seq fastq files (27, 28) are available in ENA database (https://www.ebi.ac.uk/ena/browser/home) with the following run accession numbers: ERR637895 (NF-YA), ERR637890 (NF-YB), ERR637883 (NF-YC), ERR2732089 (Sp2), ERR2732087 (PBX1), ERR2732088 (PREP1). Protein sequences were retrieved from UniProt database (https://www.uniprot.org/). The sequences used for structural predictions and EMSA oligonucleotides are listed in **Supplementary Table 1**. AlphaFold models reported in the study are listed in **Supplementary Table 2** and have been deposited in ModelArchive (https://modelarchive.org/). The models will be made publicly available upon publication. Cross-links and mutants mapped in **Fig. 4** were retrieved from the supplementary material of a previous publication (13) and are listed in **Supplementary Tables 3-4**.

