## Supplementary Figures for "Cooperative reading of DECA-CCAAT composite element by the TALE/NF-Y/Sp2 transcription factors"

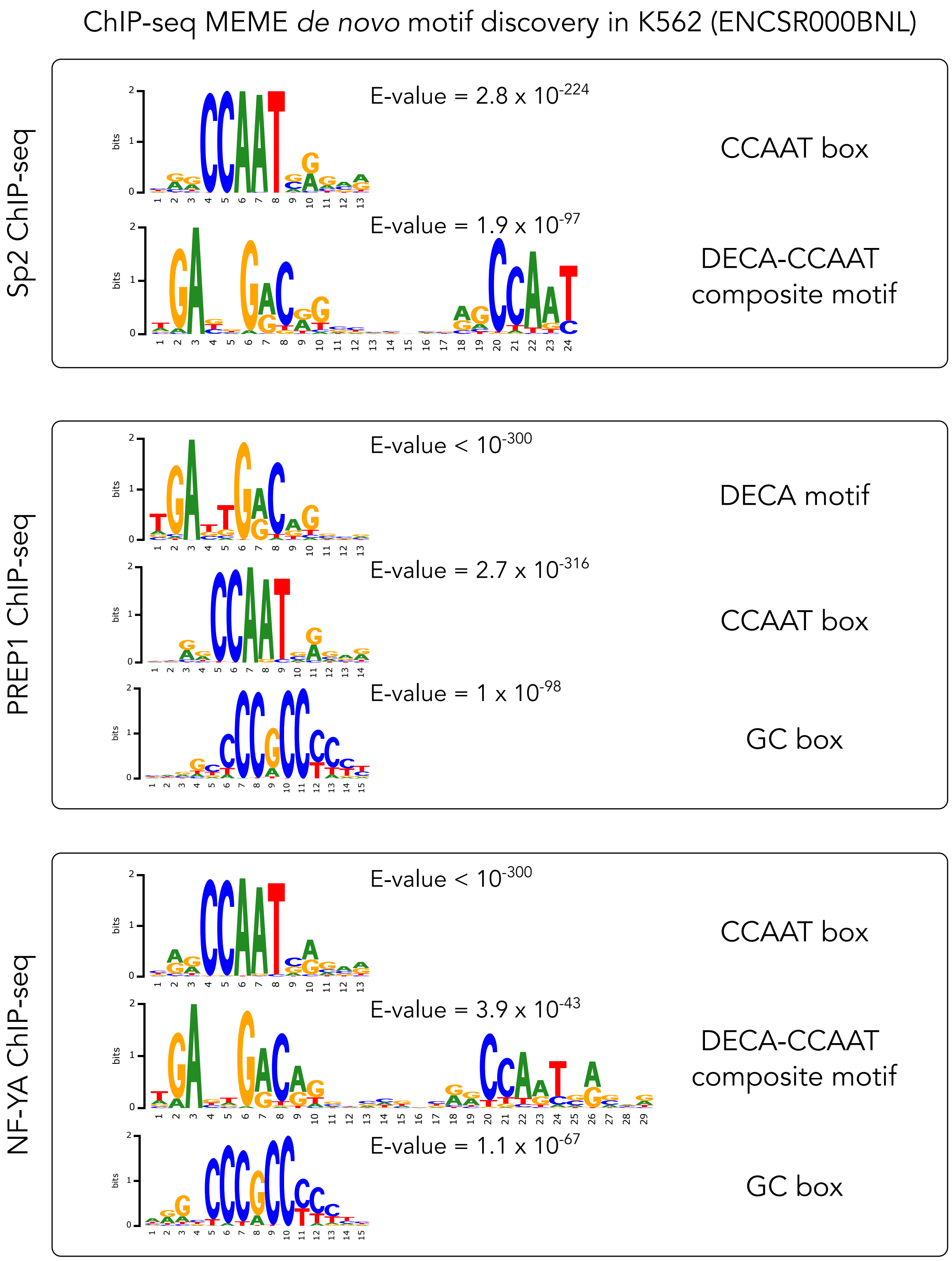


**Supplementary Figure 1. Sp2, PREP1 and NF-YA motif enrichment analysis.** Sequence logos for motifs found enriched by MEME in ChIP-seq experiments of Sp2, PREP1 and NF-YA. Data are from human K562 cells (ENCSR000BNL), as reported in Factorbook. For each logo, the corresponding E-value and motif name are indicated.

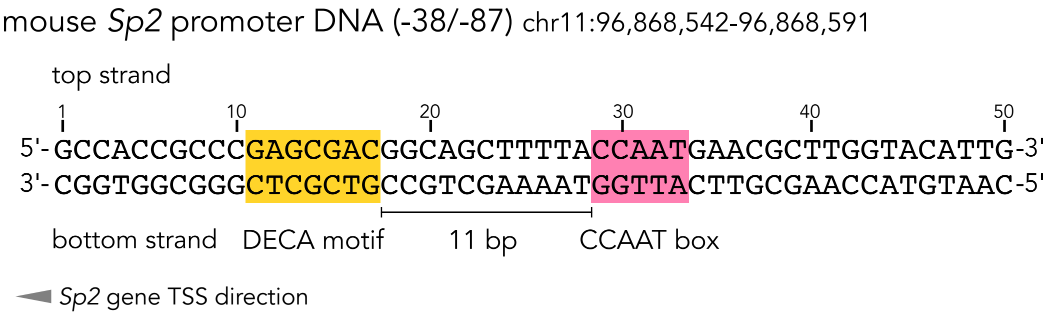


**Supplementary Figure 2. DNA sequence of mouse *Sp2* promoter used in AF modelling.** The location of the core DECA (GAnnGAC) and CCAAT motifs is indicated in yellow and pink, respectively.

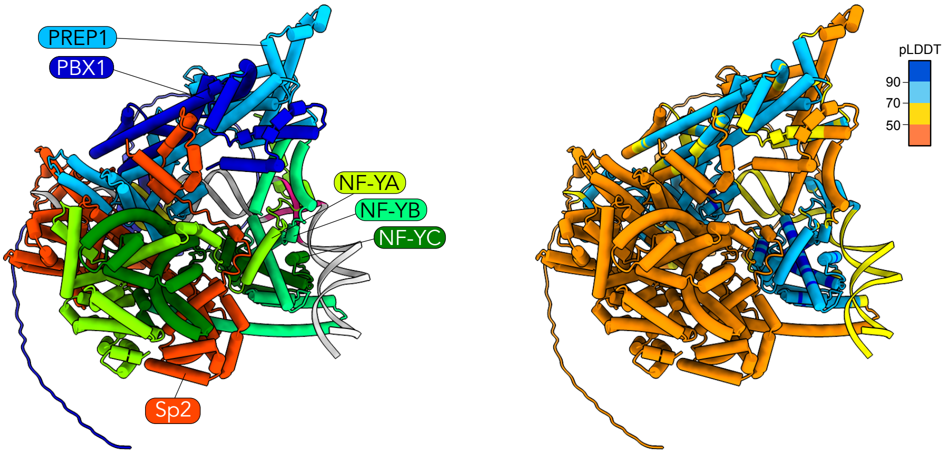


**Supplementary Figure 3. The full-length AlphaFold 3 model is subject to spurious structural order (hallucinations).** The PBX1/PREP1/NF-Y/Sp2 complex on DECA-CCAAT site was modelled using full-length proteins using AlphaFold 3. It resulted in evident forms of hallucination involving the intrinsically disordered regions of each protein, modelled as compact low-confidence $\alpha$-helical domains.

### Supplementary Figure 4

A

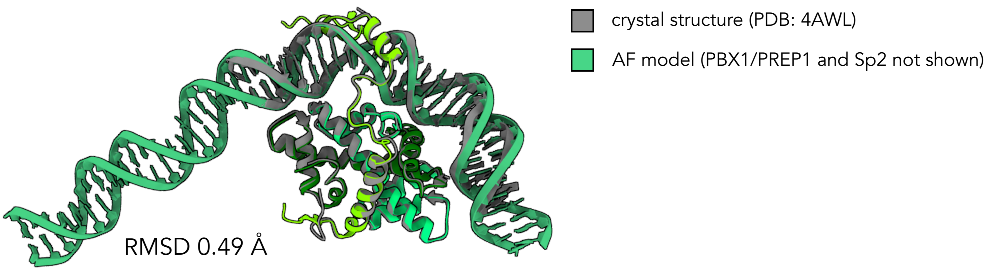


B

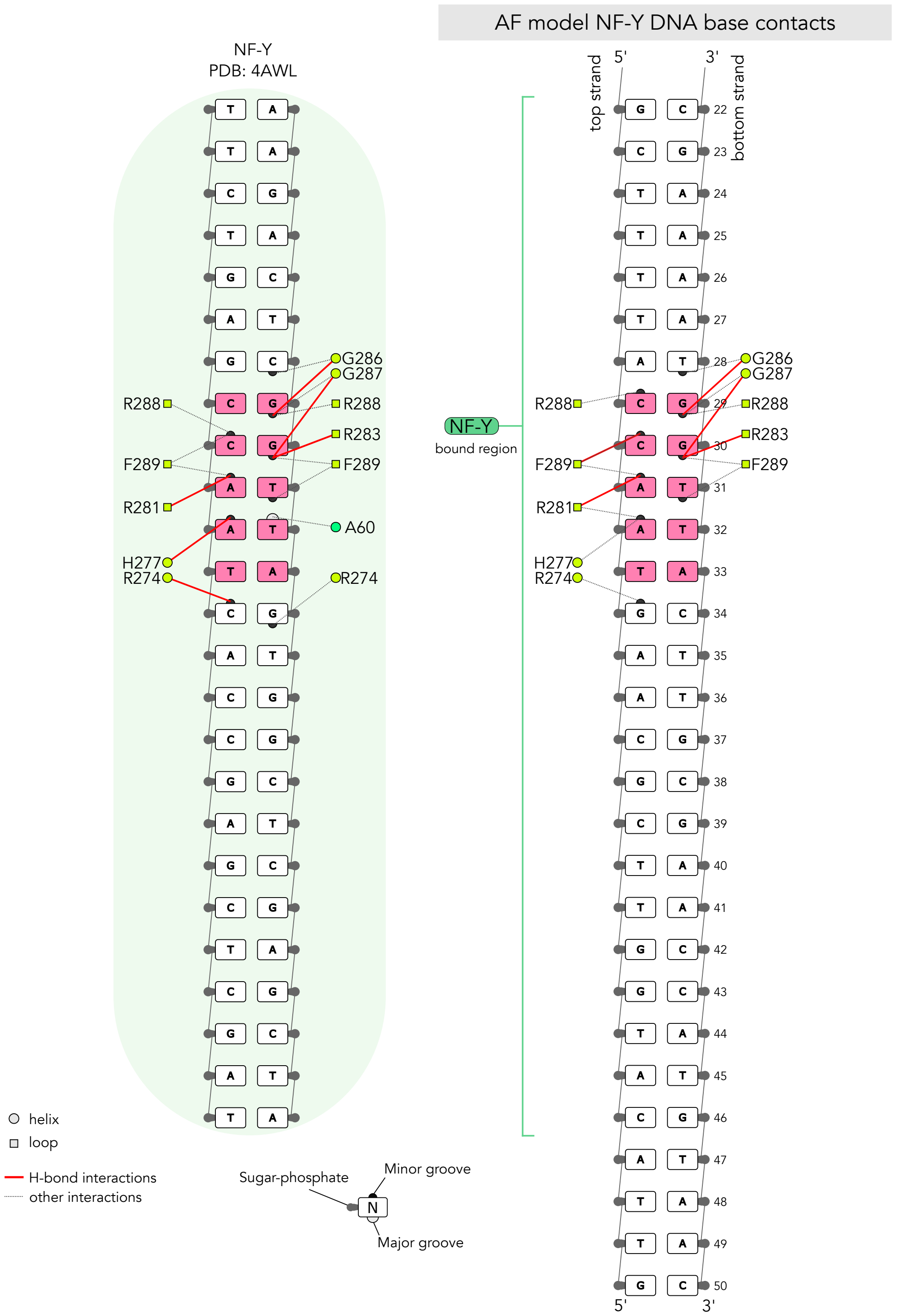

**Supplementary Figure 4. Comparison of NF-Y/CCAAT protein-nucleobase contacts in the crystal structure and AlphaFold 3 model. A**. NF-Y crystal structure (PDB: 4AWL) aligned to the model shown in Fig. 2 generated by AlphaFold 3 (only NF-Y and DNA are shown for clarity; models are aligned on NF-YB chain). **B**. Schematic representation of direct protein contacts with DNA nucleobases in the NF-Y crystal structure (left panel) *versus* the complex modelled by AlphaFold 3 on mouse *Sp2* promoter sequence. Contacts with DNA sugar-phosphate backbone are omitted. The CCAAT pentanucleotide is highlighted in pink. All contacts in the model are mediated by the NF-YA subunit (residues colored in lime green).

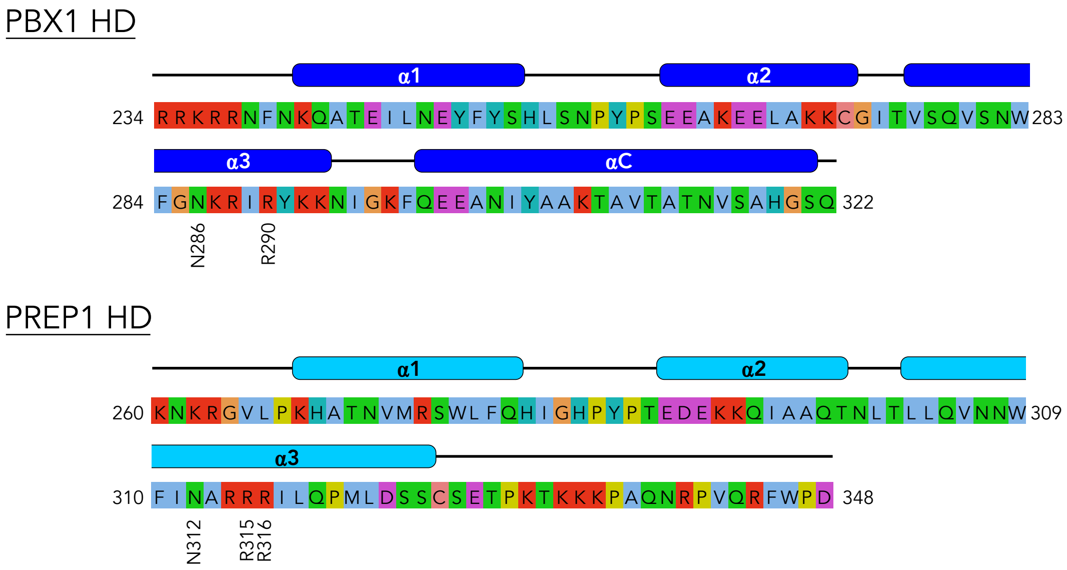


**Supplementary Figure 5. PBX1 and PREP1 homeodomain (HD) sequence.** Protein sequence of PBX1 HD (upper panel) and PREP1 HD (bottom panel). The sequence is coloured according to the Clustal colour scheme. The α-helices predicted with high confidence (pLDDT > 70) by AlphaFold are indicated above the alignment. Residues of helix 3 recognizing the GAnnGAC DECA core motif nucleobases in the AF model are indicated.

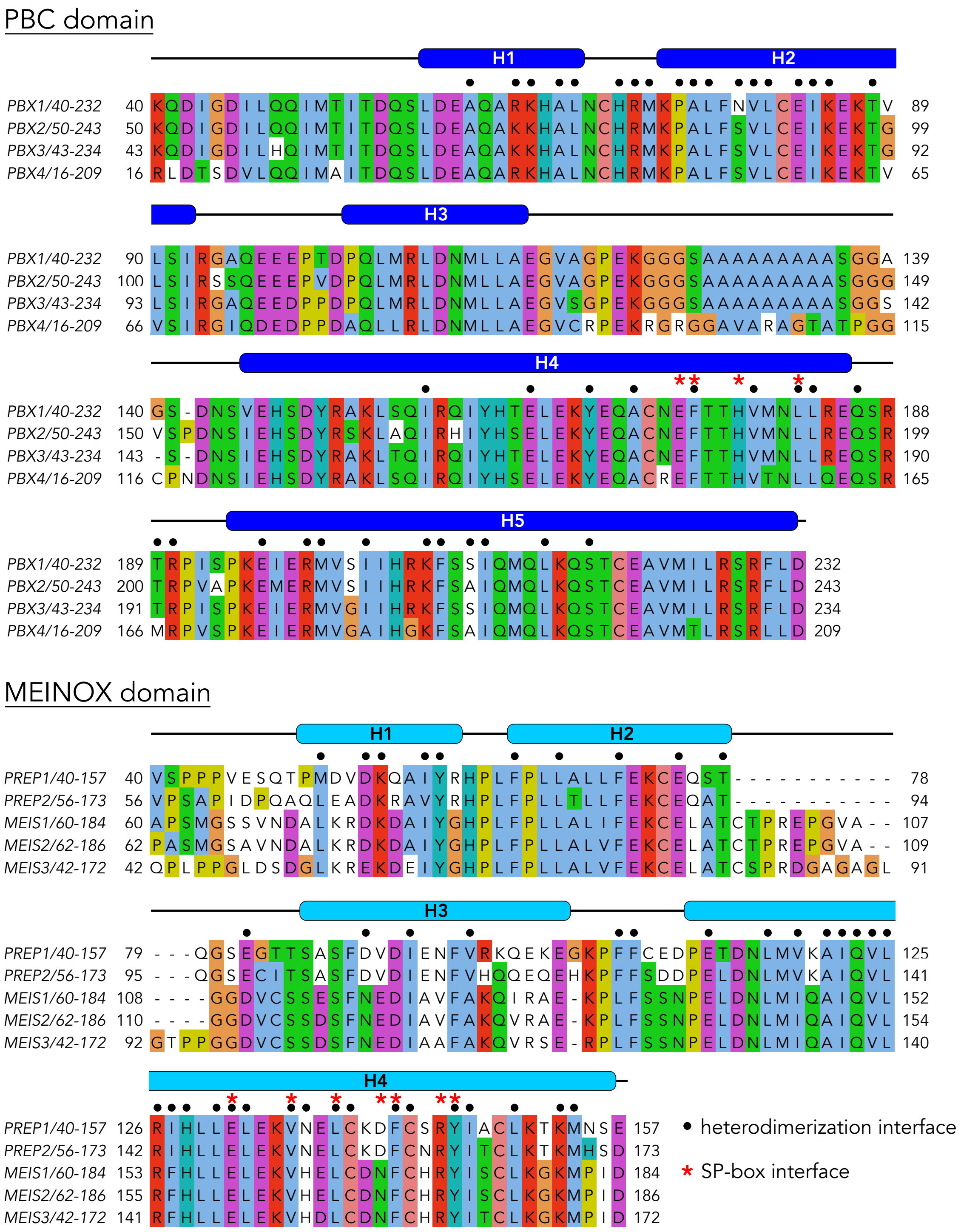


**Supplementary Figure 6. Alignment of the PBC and MEINOX heterodimerization domains.** Alignment of the heterodimerization domains of human PBX paralogs (upper panel, PBC domain) and PREP/MEIS paralogs (bottom panel, MEINOX domain). The alignment is coloured according to the Clustal colour scheme. The α-helices (H) predicted with high confidence (pLDDT > 70) by AlphaFold are indicated above the alignment. Residues positioned at the PBX1/PREP1 heterodimerization interface or at the interface with Sp2 (SP-box) in the AlphaFold model are indicated with black circles and red stars, respectively.

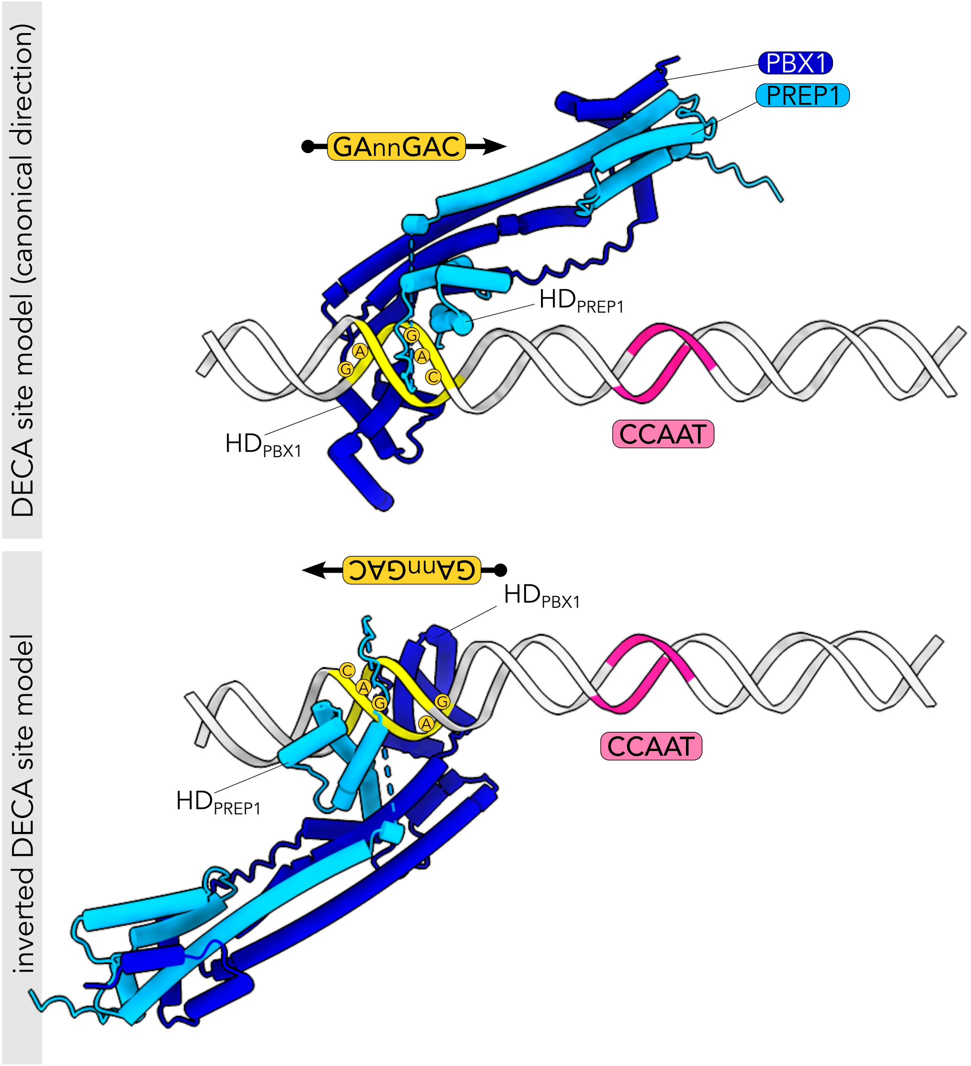


**Supplementary Figure 7. DECA motif orientation impacts on the direction of the PBC-MEINOX heterodimerization module.** The PBX1/PREP1 heterodimer was modelled on the mouse *Sp2* promoter DNA. The DECA site (in yellow) was placed either in the canonical (top panel) or reverse orientation (bottom panel). The PBX1/PREP1 heterodimer reorients according to DECA motif direction. The dashed line indicates PREP1 protein regions connected by a disordered loop (not shown for clarity).

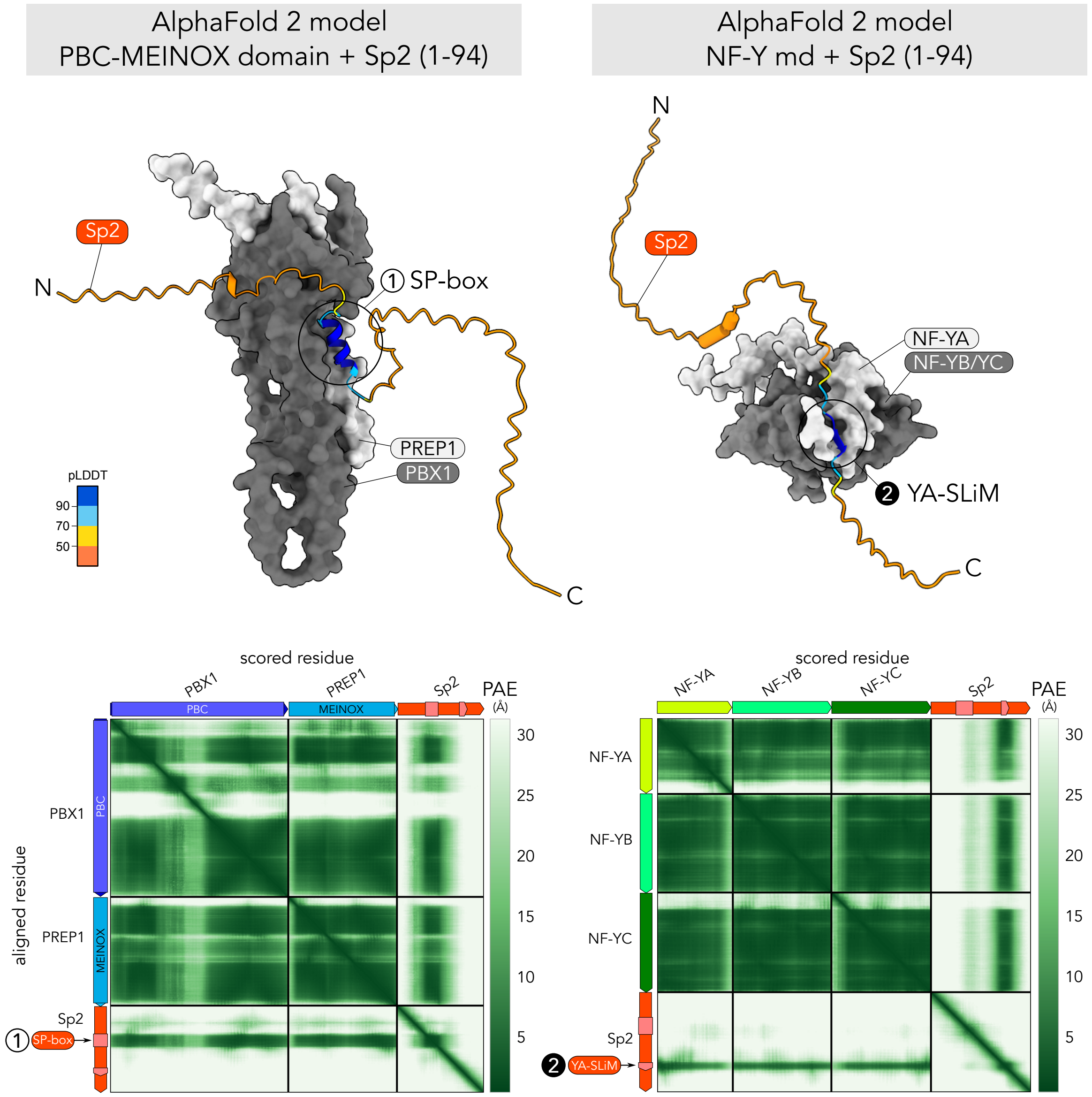


**Supplementary Figure 8. Modelling of the two Sp2 anchor points with AlphaFold2.** Representative structural models of Sp2 (aa 1-94) with PBX1-PREP1 heterodimerization module (left panels) and NF-Y (right panels) modelled using AlphaFold2-multimer. Sp2 is coloured according to the pLDDT confidence score, while the interacting partners are coloured in shades of gray as surface representations. The bottom panels show the predicted alignment error (PAE) heatmaps for each model. The positions of anchor points #1 and #2 are indicated.

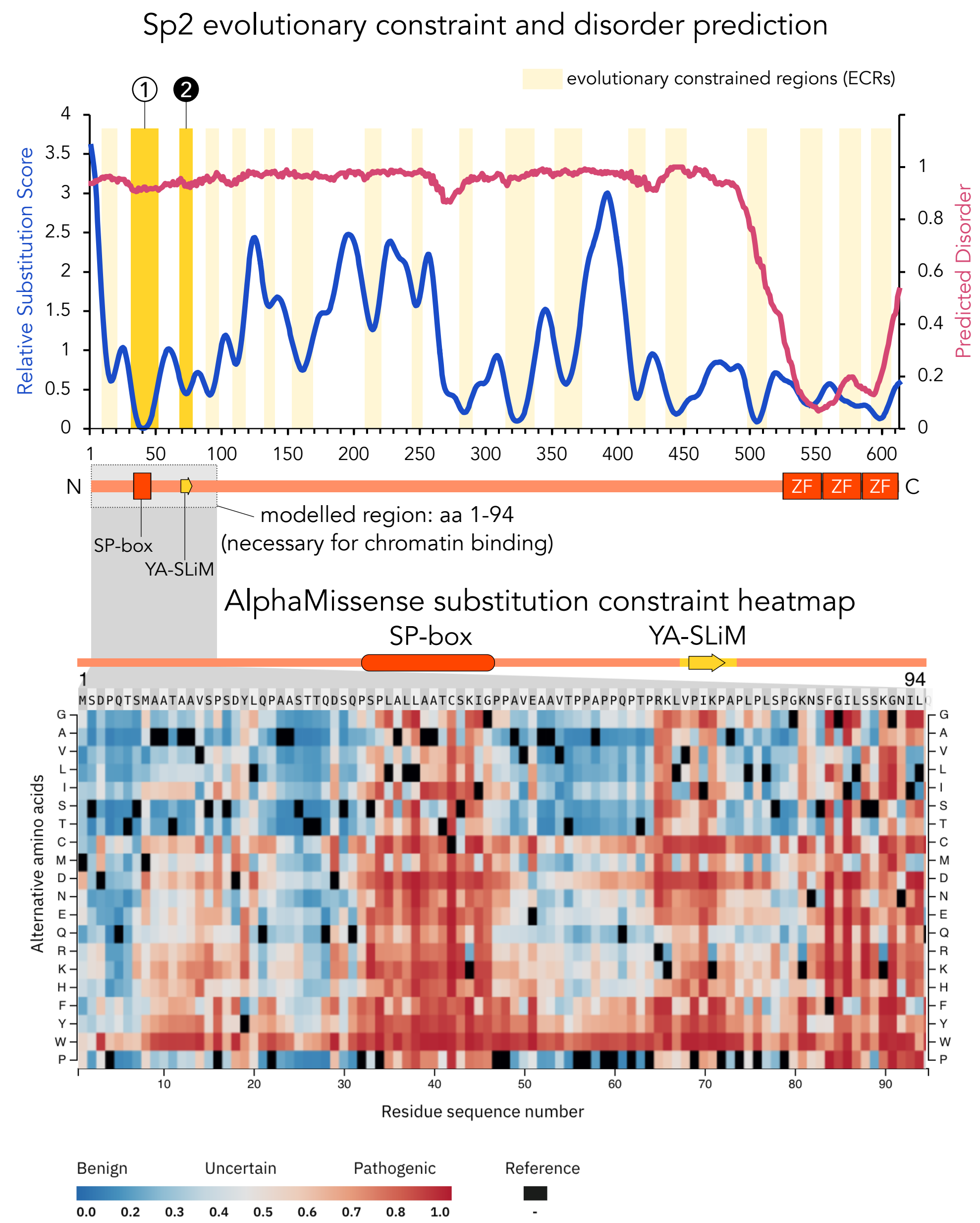


**Supplementary Figure 9. Sp2 disorder prediction and evolutionary constrained regions.** (Top panel) Prediction of intrinsic protein disorder for Sp2 (from Metapredict) is plotted along with the smoothed relative substitution score obtained from AMINODE. Regions with local substitution minima across species are tagged as evolutionary constrained regions (ECRs, shaded in yellow). Anchor points #1 (SP-box) and #2 (YA-SLiM) overlap with the second and third ECR, respectively.

(Bottom panel) AlphaMissense pathogenicity heatmap of the Sp2 region used for modelling (aa 1-94). Red cells indicate substitutions predicted to be deleterious (likely pathogenic). Red regions can be interpreted as evolutionarily constrained and functionally important.

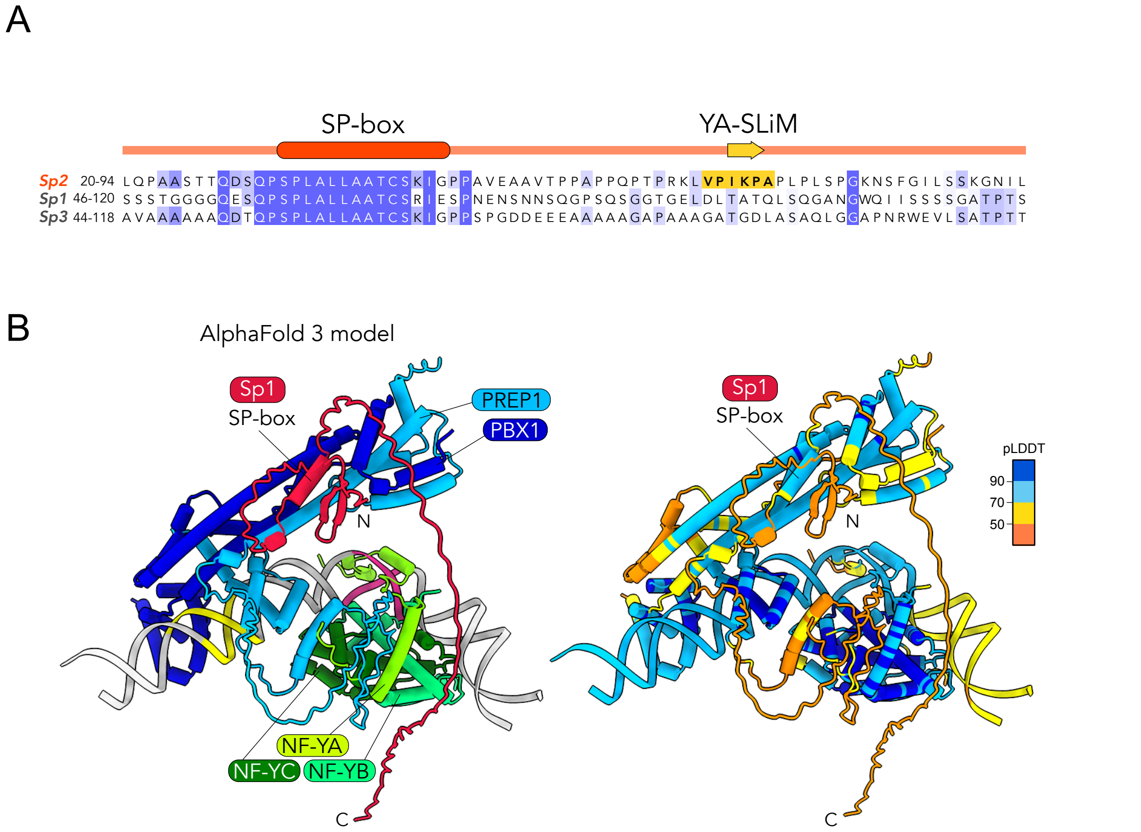


**Supplementary Figure 10. YA-SLiM is not present in other Sp-family members.**
**A**. Alignment of Sp2 with the related Sp1 and Sp3 human proteins. The alignment is coloured according to the level of sequence identity. Above the alignment the position of the SP-box and YA-SLiM is indicated. YA-SLiM is highlighted in yellow. **B**. AlphaFold 3 model of the PBX1/PREP1/NF-Y/Sp1 complex on DNA. To generate the model, Sp2 was substituted with Sp1 (aa 1-120) in the input sequences used to build the model shown in Fig. 2. The right panel shows the same model coloured according to the pLDDT confidence score.

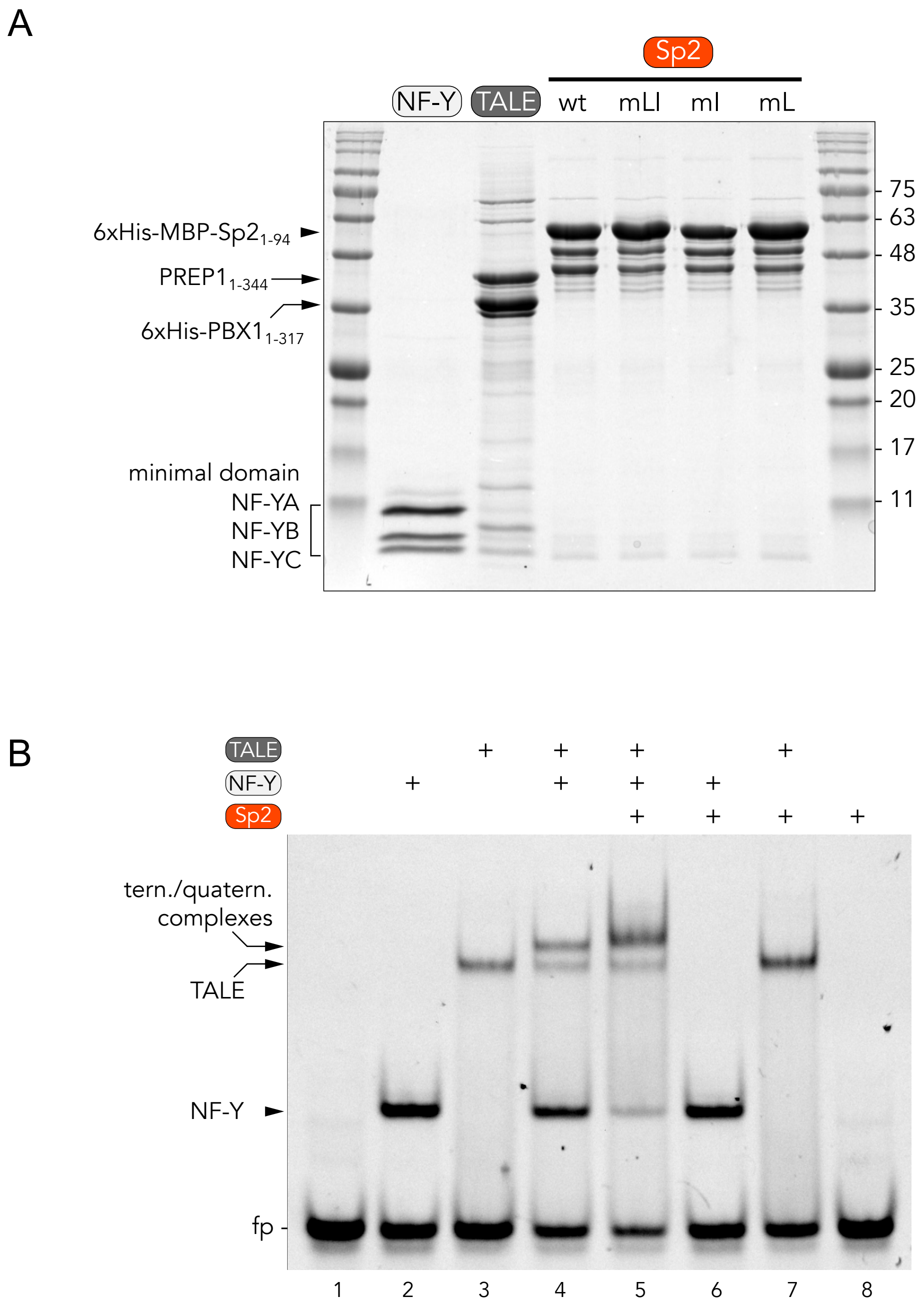


**Supplementary Figure 11. Recombinant proteins and EMSA controls.**
**A**. Coomassie Brilliant Blue stained SDS-PAGE gel of purified recombinant proteins used in EMSA experiments (3 µg NF-Y, 5 µg TALE, 3 µg Sp2 were loaded on the gel). **B**. EMSA experiment with different combinations of the recombinant proteins shown in A. Proteins concentration in the assay is the following: 40 nM NF-Y, 60 nM TALE, 320 nM Sp2-wt. DNA probe concentration is 20 nM. The different DNA-bound complexes and the free probe (fp) are indicated on the left.

SP-box alignment

```
>Sp2 (33-46)          SPLALLAATCSKIG
>Sp1 (59-72)          SPLALLAATCSRIE
>Sp3 (57-70)          SPLALLAATCSKIG
>Sp4 (46-59)          SPLALLAATCSKIG
>Sp5 (31-44)          SPLALLAATCSRIG
>Sp6 ()               --------------
>Sp7 (15-28)          SPLAMLTAACSKFG
>Sp8 (15-28)          TPLAMLAATCNKIG
>Sp9 (15-28)          TPLAMLAATCNKIG

>ZNF503 (111-124)     SPLALLAQTCSQIG
>ZNF703 (87-100)      SPLALLAQTCSQIG
```

**Supplementary Figure 12. SP-box in Sp-family members and ZNF503/ZNF703.** Amino acid sequences of the SP-box motif are aligned and the correspondent sequence range is indicated on the left. Only positions identical to the Sp2 sequence are colored (according to the aminoacid type).
